# A Lattice Model on the Rate of DNA Hybridization

**DOI:** 10.1101/2021.12.22.473940

**Authors:** R. Murugan

## Abstract

We develop a lattice model on the rate of hybridization of the complementary single-stranded DNAs (c-ssDNAs). Upon translational diffusion mediated collisions, c-ssDNAs interpenetrate each other to form correct (cc), incorrect (icc) and trap-correct contacts (tcc) inside the reaction volume. Correct contacts are those with exact registry matches which leads to nucleation and zipping. Incorrect contacts are the mismatch contacts which are less stable compared to tcc which can occur in the repetitive c-ssDNAs. Although tcc possess registry match within the repeating sequences, they are incorrect contacts in the view of the whole c-ssDNAs. The nucleation rate (*k_N_*) is directly proportional to the collision rate and the average number of correct-contacts (<*n_cc_*>) formed when both the c-ssDNAs interpenetrate each other. Detailed lattice model simulations suggest that 〈*n_cc_*〉 ∝ *L*/*V* where *L* is the length of c-ssDNAs and *V* is the reaction volume. Further numerical analysis revealed the scaling for the average radius of gyration of c-ssDNAs (R_g_) with their length as 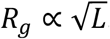. Since the reaction space will be approximately a sphere with radius equals to 2*R_g_* and *V* ∝ *L*^3/2^, one obtains 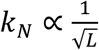. When c-ssDNAs are nonrepetitive, then the overall renaturation rate becomes as *k_R_* ∝ *k_N_L* and one finally obtains 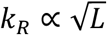 in line with the experimental observations. When c-ssDNAs are repetitive with a complexity of *c*, then earlier models suggested the scaling 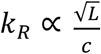 which breaks down at *c* = *L*. This clearly suggested the existence of at least two different pathways of renaturation in case of repetitive c-ssDNAs viz. via incorrect contacts and trap correct contacts. The trap correct contacts can lead to the formation of partial duplexes which can keep the complementary strands in the close vicinity for a prolonged timescale. This is essential for the extended 1D slithering, inchworm movements and internal displacement mechanisms which can accelerate the searching for the correct contacts. Clearly, the extent of slithering dynamics will be inversely proportional to the complexity. When the complexity is close to the length of c-ssDNAs, then the pathway via incorrect contacts will dominate. When the complexity is much lesser than the length of c-ssDNA, then pathway via trap correct contacts would be the dominating one.

**PACS:** 87.10.-e; 87.14.gk; 82.39.Pj; 87.15.R-

## 1. Introduction

The reversible unwind-rewind property of the double stranded helical structure of DNA is critical for its various biological functions [1,2]. The double helical structure of DNA (**dsDNA**) that is stabilized by the hydrogen bonding network and hydrophobic base-stacking at the core [1]. Denaturation is the process of melting of dsDNA into the corresponding complementary single strands [2]. These single strands of DNA (**ssDNA**) zip back spontaneously to form the original dsDNA upon removal of the denaturant which is known as renaturation or hybridization [2–4]. Transcription, translation and replication of the gnomic DNA and several *in vitro* laboratory techniques are based on the denaturation-renaturation property of dsDNA [5]. Clear understanding on the mechanism of DNA hybridization in solution is important to design efficient primers for polymerase chain reaction (PCR), design of oligonucleotide probes for microarray chips, design and construction of versatile nano-structures over single stranded DNA scaffolds using DNA origami method [6,7] and various DNA fingerprinting technologies [5]. Detailed understanding of the mechanism of hybridization of ssDNAs at the microscopic level is one of the contemporary issues in the biological physics field.

Several models on DNA renaturation have been developed and experimentally tested [2,8–17]. There are two different views viz. one-step and two-step models. Hybridization of the complementary ssDNAs (**c-ssDNAs**) was initially described as a one-step diffusion controlled bimolecular collision process as in **Scheme I** of **Fig. 1A** [4,18–20]. Although this scheme was simple enough to capture most of the underlying dynamics [8,14,17,19], it could not explain several other polymer physics related scaling relationships. Particularly, **Scheme I** predicted a linear scaling of the bimolecular collision rate with respect to the length of c-ssDNAs. Whereas, the experimental data revealed approximately a square root type scaling [8,14]. Further experiments revealed an inverse scaling of the bimolecular collision rate with the sequence complexity of ssDNA.

**FIGURE 1.**
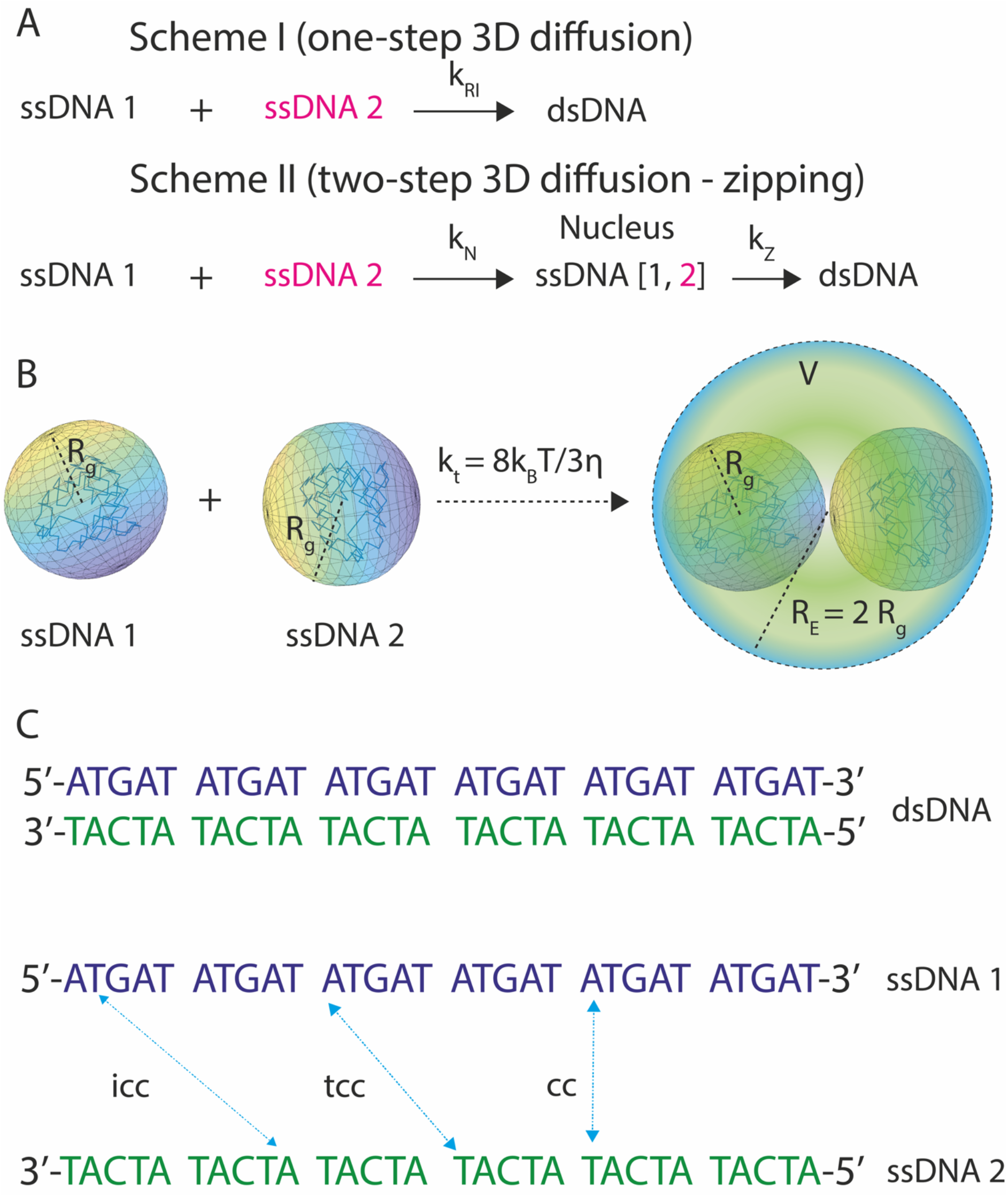
Models on DNA renaturation. **A**. one-step, two-step models. In one-step model described in **Scheme I**, the complementary strands forms duplex with a second order rate *k_RI_* directly by diffusion controlled 3D collisions. In the two-step model described in **Scheme II**, the renaturation progress by the formation of stable nucleus with a rate *k_N_* via 3D diffusion mediated collisions and interpenetrations of c-ssDNAs and then subsequently by zipping with a rate *k_Z_*. **B**. The complementary ssDNA strands can be thought as spherical-shaped and loosely-packed nucleotide clusters with average radius of gyration *R_g_*. Upon translational diffusion, they arrive inside the reaction volume V with a rate *k_t_* and interpenetrate each other. Here the reaction space is assumed to be a spherical one with radius 2*R_g_*. When the complementary strands are confined inside *V*, there may be contacts between them. **C**. When the contact occurs with exact registry match, then it is a correct-contact that can lead to nucleation and zipping. When contacts occur between nonidentical registers, then they are all incorrect contacts. When c-ssDNAs contain repeats, then there may be contacts between identical registries of repeats placed at two nonidentical locations. These are trap correct-contacts which can lead to the formation of partial duplexes with single strand overhangs. In the given example, there are six repeats. When the position 1 of first repeat interact with position 1 of second repeat, then it is a tcc. Sequence complexity is defined as the number of bases in an unique sequence. For example, the given sequence contains the repeats of 5 bases. Therefore, its complexity is *c* = 5 where the total length is *L* = 30.

Wetmur and Davidson [8] suggested a detailed two-step model with nucleation and zipping as in **Scheme II** of **Fig. 1A**. In their model, the overall renaturation rate was directly proportional to the product of nucleation rate and length of c-ssDNAs and inversely proportional to the sequence complexity. Here, the nucleation rate scales with the length of c-ssDNA in an inverse square root manner. As a result, the overall renaturation rate scales with the length of ssDNA in a square root manner. They further argued that the renaturation was not a diffusion controlled process, and the inverse square root scaling of the nucleation rate with the length of c-ssDNA must be due to either the excluded volume effects associated with the intra-strand dynamics or steric hindrance associated with the interpenetration of c-ssDNAs that is essential for the nucleation [8]. However, this argument did not explain the inverse scaling of the renaturation rate with the viscosity of the medium. To overcome this issue, they further assumed a diffusion mediated growth of the nucleus with a rate that is inversely related with the viscosity coefficient of the medium [21]. This assumption ensured the inverse scaling of the overall renaturation rate with the viscosity of the medium.

The c-ssDNAs are random coils with average radius of gyration *R_g_* (**Fig. 1B**). At coarse grained level, one can think both the complementary strands as loosely packed spherical clusters of nucleotide monomers with radius *R_g_* performing translational diffusion. The base clusters of c-ssDNAs interpenetrate each other upon collision and subsequently nucleation occurs inside the reaction volume. Clearly, there are two distinct dynamical regimes in the process of renaturation viz. 1) translational diffusion mediated collision of base clusters of c-ssDNAs and 2) their interpenetration. Translational diffusion brings both the strands within the reaction radius. Reaction radius will be the sum of the radius of gyration of c-ssDNAs. Subsequently, these c-ssDNA base clusters interpenetrate each other within the reaction volume to achieve the nucleation and zipping. Therefore, the translational diffusion component cannot be ignored here. Further, the expression for the reaction rate corresponding to the two-step model (Eq. 3 in Ref. [8]) will be inconsistent whenever the sequence complexity has the same magnitude as that of the length of c-ssDNAs. Under such scenario, the overall renaturation rate would inversely scale with the length of c-ssDNAs in a square root manner that is inconsistent with the experimental observations [8]. Here the *sequence complexity* is defined as the length of DNA with unique nucleotide sequence pattern (**Fig. 1C**). Several theoretical and computational models [13,22–28] were developed recently to explain the observed scaling behaviours of the overall second order renaturation rate on the size of c-ssDNAs, temperature, ionic strength and viscosity of the reaction medium.

The nucleation step can be modelled as Kramer’s escape problem over a free energy barrier [24]. However, nature of the reaction coordinate and the entropic component of potential energy barrier associated with the renaturation process is unclear. According to the recently proposed three-step model [15] (**Scheme III** in **Fig. 2**), the renaturation process comprises of (**a**) formation of nonspecific contact (**b**) nucleation or correct contact formation and (**c**) zipping. In the first step, c-ssDNAs perform three-dimensional collisions which result in the formation of Watson-Crick (WC) base pairs at random nonspecific contacts. Such nonspecific contacts randomly translocate along c-ssDNAs either via thermally driven one-dimensional (1D) slithering, inchworm movements and internal displacement [25] mechanisms until finding the correct-contact to initiate the nucleation process. Nucleation will be followed by spontaneous zippering of c-ssDNAs. In this model, the square root dependency of the renaturation rate on the length of c-ssDNAs mainly originates from the non-specific contact formation step. However, it is still not clear whether the mode of nucleation is via 1D or 3D diffusion. It is also not clear how the inverse scaling on the sequence complexity arises in case of repetitive c-ssDNAs. Further, the retardation effects of the repetitive DNA sequences were not considered in Scheme III of Ref.[15].

**FIGURE 2.**
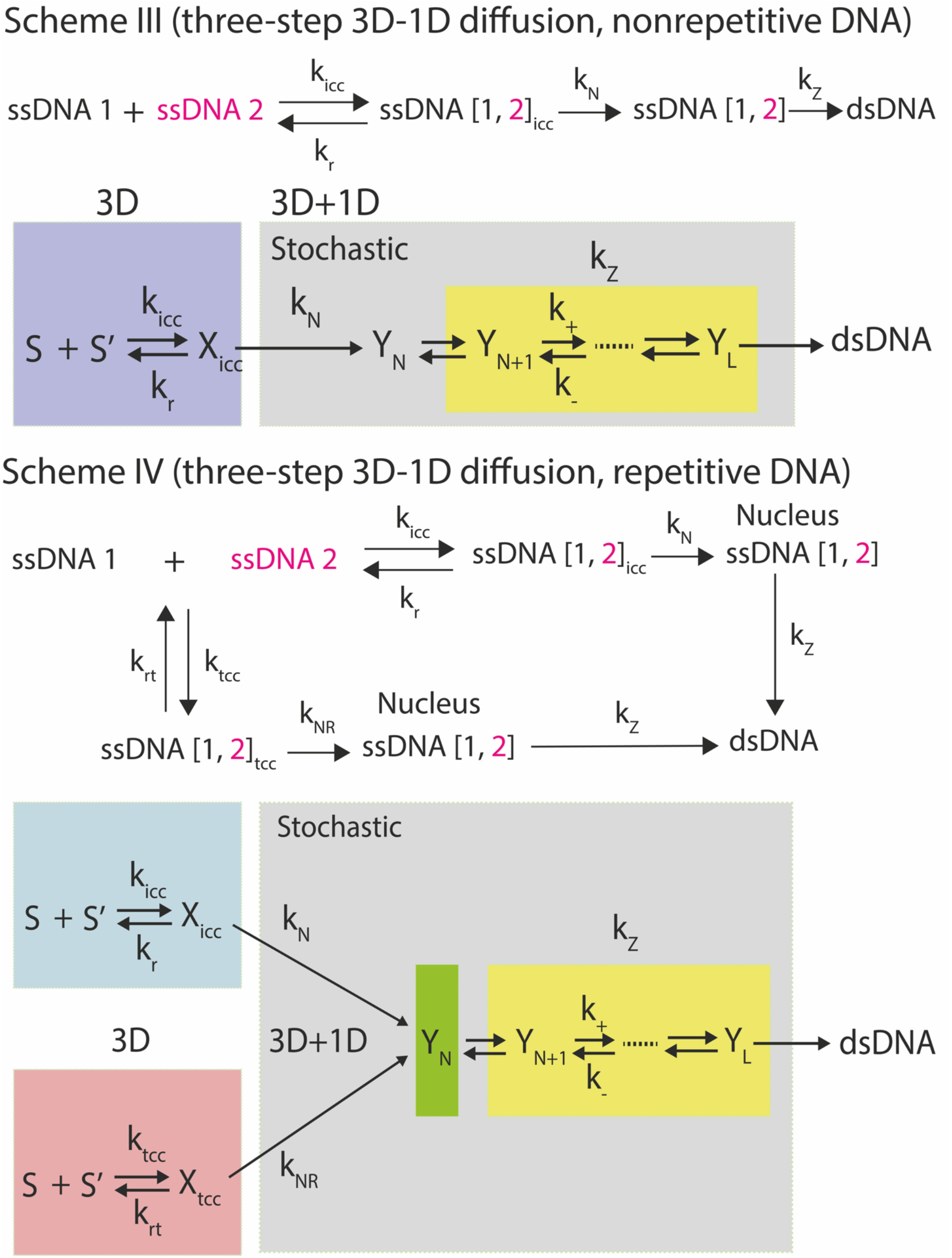
Three step models on DNA renaturation. Here S and S’ represents ssDNA1 and ssDNA2 respectively, X_icc_ = ssDNA [1,2]_icc_, X_tcc_ = ssDNA [1,2]_tcc_, *Y_N_* represents the nucleus with *N* nucleotides. In the three-step model given by **Scheme III**, c-ssDNAs renature via a combination of 3D and 1D diffusions. Here *k_tcc_* and *k_icc_* are the number of incorrect and trap-correct contacts formed per second, *k_r_* and *k_rt_* are respective dissociation rate constants, *k_N_* and *k_NR_* are the respective nucleation rates and *k_+_* an *k*_−_ are the forward and reverse microscopic rate constants associated with the zipping steps. In Scheme III, 3D collisions result in the formation of incorrect contacts which bring the strands into the close vicinity. Nucleation occurs via various 1D diffusion dynamics such as slithering, inchworm movements and internal displacement mechanisms. When c-ssDNAs are repetitive, then there is always chances for the formation of partial duplexes with single strand overhangs. This will in turn retain the c-ssDNAs in the close vicinity for prolonged amount of time that results in *k_r_* > *k_rt_* and long slithering lengths. This will be a parallel pathway along with **Scheme III** as demonstrated in **Scheme IV** in case of repetitive c-ssDNAs.

Understanding the role of the conformational state of DNA on the rate of hybridization is critical to unravel the underlying mechanism. To understand the equilibrium thermodynamic properties of the DNA hybridization, several course grained lattice models were developed and investigated using Monte Carlo simulation methods [28]. However, simulation of such systems with detailed base-paring and base-stacking interactions were limited to very short DNA sequences [25,28]. Recently, Qu, et.al. [29] have simulated up to around 85 nucleotides using the oxDNA GPU-enabled standalone code. Their simulated time-dependent system energy profile exhibited a four-state hybridization mechanism in line with earlier theoretical models [15]. Using similar codes, Snodin, et.al. [30] simulated the hybridization kinetics of DNA origami for a system of 384 nucleotides. Such studies are useful to obtain the equilibrium thermodynamic properties rather than the nonequilibrium kinetic scaling aspects. In this context, short strands have been studied using non-equilibrium methods for enhanced forward-flux sampling at much reduced computational requirements. Particularly, Schreck, et.al. [31] have studied 25 nucleotide oxDNA, Gravina, et.al. [32] have studied 42 nucleotide 3SPN.2 DNA, Jones, et.al. [33] have studied a DNA with 10 nucleotides AT repeats. Apart from these, Sidky, et al. [34] have created a Software Suite for Advanced General Ensemble Simulation (SSAGES) which can use the parallel computational workflows for forward-flux sampling. Clearly, all these computational methods assume that the interacting c-ssDNAs are already arrived at the reaction volume through translational diffusion. Whereas, the kinetic scaling laws with respect to varying lengths of DNA, sequence complexity and viscosity of the reaction medium associated with the hybridization are mainly dependent on the translational diffusion of c-ssDNAs along with their interpenetration dynamics rather than the fine details of base-pairing and base-stacking interactions which are main focus of all the coarse grained Monte Carlo simulation methods. Further, though such computational studies revealed several interesting qualitative aspects of hybridization [29,33], derivation of scaling laws through a computational route requires the simulation of very large lengths of ssDNAs with different levels of sequence complexities that still limits the application of molecular dynamics methods. In this background, we model c-ssDNAs as self-avoiding random walks (**SARW**) confined in a cubic lattice box which represents the reaction volume under *in vitro* conditions. Using this lattice model along with the observations from the computational studies [29,33], we will unravel the origin of various kinetic scaling relationships associated with the renaturation dynamics in detail.

## 2. Theoretical Methods

### 2.1. Translational diffusion of c-ssDNAs

We model c-ssDNAs (we denote them as ssDNA 1 and 2 as in **Fig. 1A**) as loosely packed and approximately spherical shaped nucleotide clusters with a radius of gyration *R_g_*. The 3D collisions between these c-ssDNAs are catered via their translational diffusion. When the length of both these strands are equal to *L*, then one can approximately assume equal radius of gyration *R_g_* for both the strands. In this situation, the maximum possible steady state Smolochowski rate of 3D diffusion controlled collisions between these c-ssDNAs will be 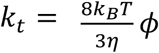 [35] (**Fig. 1B**). Here *k_B_* is the Boltzmann constant, *η* is the viscosity coefficient of the medium, *T* is the absolute temperature in degrees *K* and 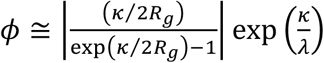 [15,36] is the multiplication factor associated with the electrostatic repulsions between the negatively charged phosphate backbones of c-ssDNAs along with the shielding effects of solvent molecules and other ions present at the DNA-DNA interface under dilute conditions. Here 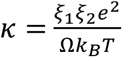 is the *Onsager radius which is defined as the distance between the colliding c-ssDNA strands at which the overall electrostatic interaction energy will be equal to that of the background thermal energy* (1 *k_B_T*), *ξ*_1_, *ξ*_2_ are the overall charge numbers on the base clusters of strand 1 and 2, λ is the thickness of the ionic shell present over the charged groups of c-ssDNAs. Here *κ* will be a positive quantity since there is a charge-charge repulsion between c-ssDNAs. Since the electrostatic interactions act only at short ranges, in general one finds that |*κ*| ≪ 2*R_g_* where the reaction radius is 2*R_g_*. As a result, 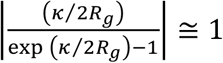 and therefore the multiplication factor 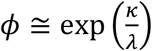 will be independent on the radius of gyration of c-ssDNAs. One also should note that *κ* < 0 in case of site-specific DNA-protein interactions since the DNA binding domains of the DNA binding proteins are usually rich in positively charged amino acids. Under such conditions, *ϕ* will be dependent on *R_g_* [37].

In the calculation of *k_t_*, we have assumed an absorbing boundary condition for the associated diffusion equation at the reaction radius that is approximately equals to 2*R_g_*. By definition, this is the rate of arrival of the base clusters of c-ssDNAs within the reaction radius. Here *k_t_* does not enumerate the number of contacts formed upon such collisions and subsequent interpenetrations. We will show in the later sections that there is always a nonzero probability for the occurrence of zero-contacts upon collision and interpenetration of c-ssDNAs. Clearly, the translational diffusion brings c-ssDNAs inside the reaction volume and there is no physical confinement of the polymer within a closed space here. The reacting c-ssDNAs enter and exit the reaction volume freely by diffusion. We will show in the later sections that this assumption is essential since the radius of gyration of c-ssDNAs will be strongly dependent on the confinement volume.

### 2.2. Interpenetration dynamics of c-ssDNAs

With this background, the nucleation rate (*k_N_*) will be directly proportional to the number of correct contacts formed between the c-ssDNAs per second. On the other hand, the average number of correct contacts formed per second (*k_cc_*) will be equal to the number of collisions between the base clusters of c-ssDNAs per second (*k_t_*, measured in M^−1^ s^−1^) multiplied by the average number of **c**orrect **c**ontacts (<*n_cc_*>) formed after each collision and interpenetration inside the reaction volume (*V*) i.e. *k_cc_* = *k_t_*〈*n_cc_*〉. As result, one obtains *k_N_* ∝ *k_t_*〈*n_cc_*〉. Only the correct-contact is the perfect registry match between the c-ssDNAs that can lead to nucleation and zipping. Apart from the correct contacts, several **i**n**c**orrect **c**ontacts (icc) can also be formed between the c-ssDNAs upon collision. When the c-ssDNAs are repetitive, then there are possibilities for the **t**rap **c**orrect **c**ontacts (tcc) as described in **Fig. 1C**. Trap correct contacts possess registry matches within the repeats but they are all incorrect contacts in the view of the entire c-ssDNAs. Here tcc can lead to partial duplexes with single strand overhangs which are actually kinetic traps in the pathway of DNA renaturation. We denote the number of cc, icc and tcc as *n_cc_*, *n_icc_*, *n_tcc_* respectively. Clearly, *n_cc_*, *n_icc_*, *n_tcc_* are all random variables since they vary from collision to collision and interpenetrations. We denote the corresponding ensemble averages as <*n_cc_*>, <*n_icc_*> and <*n_tcc_*> respectively. With these definitions, one can derive the following expressions.

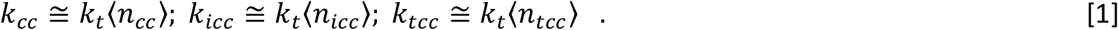

In these equations, *k_cc_, k_icc_* and *k_tcc_* are respectively the number of correct, incorrect and trap-correct contacts formed between c-ssDNAs strands per second. Clearly, the nucleation rate *k_N_* ∝ *k_cc_*. Various symbols and definitions used throughout this paper are summarized in **Table 1**. The correct-contacts can form anywhere on the entire stretch of c-ssDNAs with length equals to *L* number of nucleotides (nt). Therefore, the probability of getting a correct contact upon each collision will be *p_cc_* = 1 / *L*. This also means that the average number of correct and incorrect contacts are approximately connected via 〈*n_cc_*〉 ≅ *p_cc_*〈*n_icc_*〉. To understand various scaling behaviours of <*n_cc_*>, <*n_icc_*> and <*n_tcc_*>, we model c-ssDNAs as self-avoiding random walks (SARWs) confined in a lattice cube box that represents the reaction volume *V* of the standard bimolecular collision model. In this setting, the volume of a monomeric unit is *v* = 1 and volume occupied by the c-ssDNA strand of length *L* will be *vL*. When the complexity of ssDNA is *c* nt, then there will be at least *L* / *c* number of repeating sequences in each c-ssDNA strand. Here we assume that the respective c-ssDNA base clusters have already arrived at the reaction volume through the 3D translational diffusion with rate *k_t_*. The average number of cc, icc and tcc can be obtained by repeated generation of two independent SARWs of length *L* inside a fixed cubic lattice box with volume *V*. In each iteration, the number of cc, icc and tcc will be counted and these counts are averaged over 10^5^ SARW trajectories. Detailed stochastic simulations with fixed *L* for both the c-ssDNA strands and reaction volume *V* as given in the **Simulation Methods** section revealed the following scaling relationships.

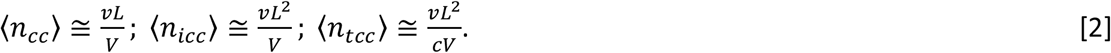

**TABLE 1.**
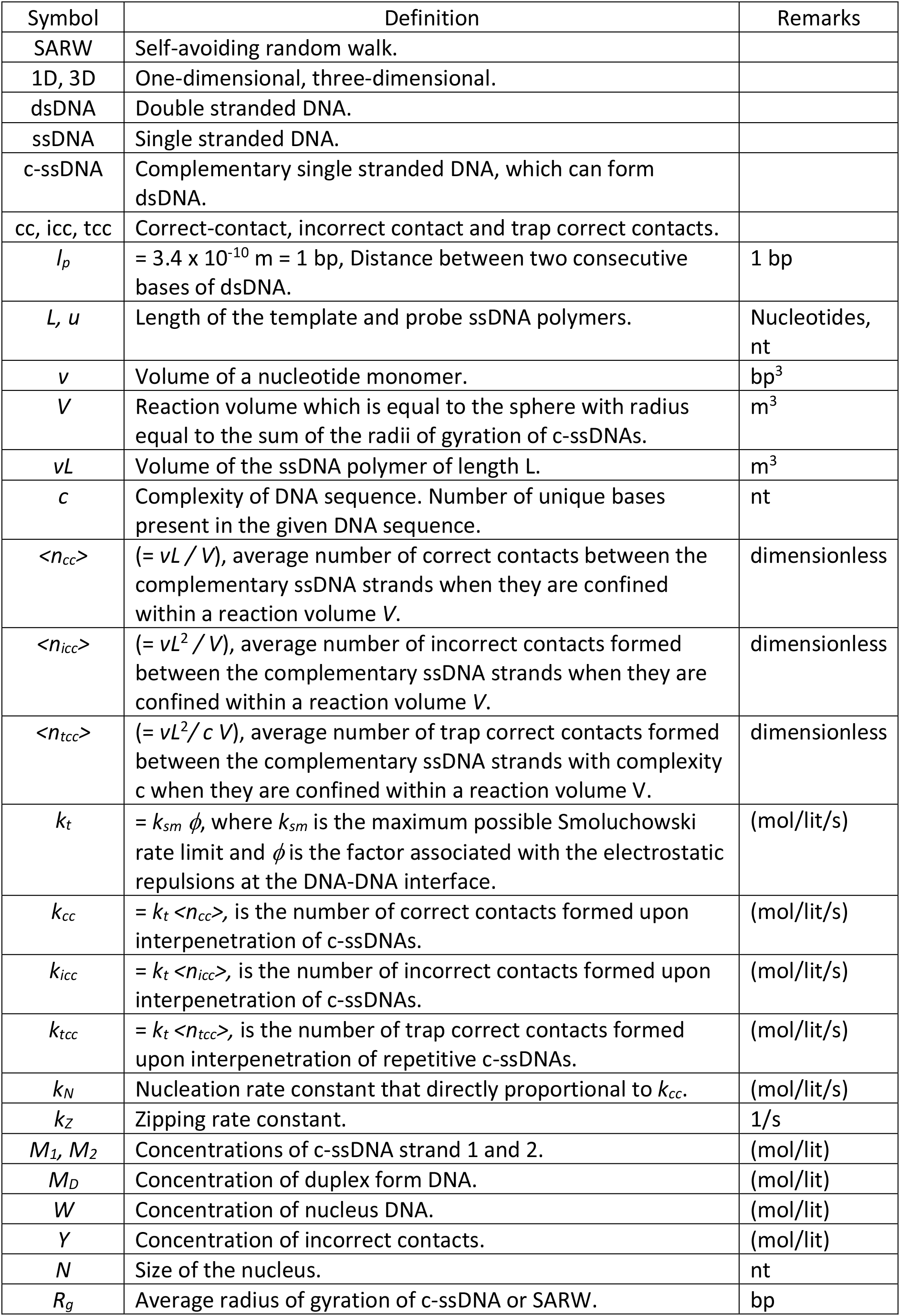
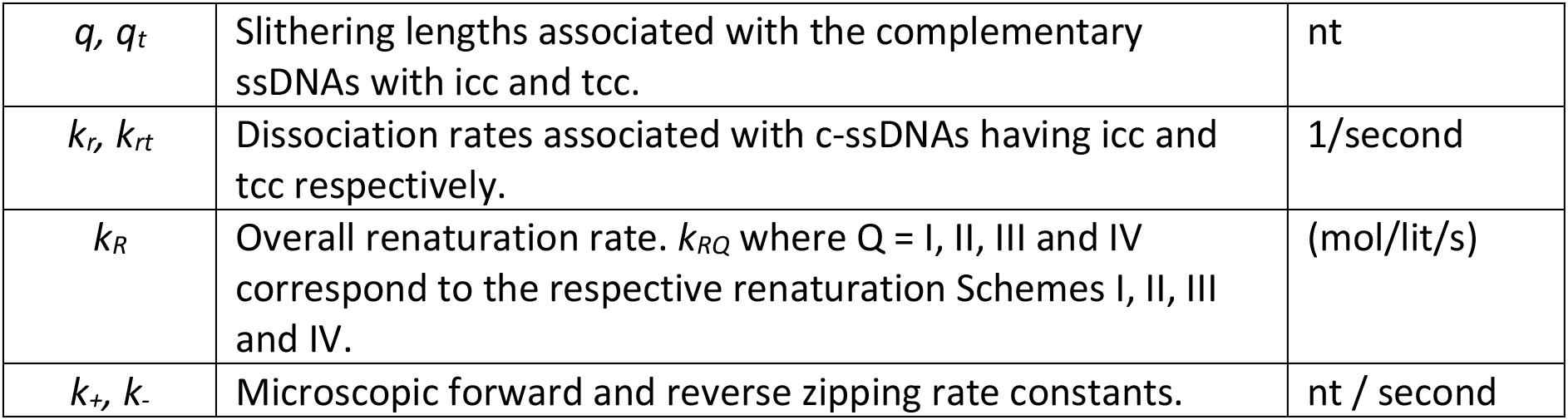
List of symbols and definitions used in the main text.

| Symbol | Definition | Remarks |
| --- | --- | --- |
| SARW | Self-avoiding random walk. |  |
| 1D, 3D | One-dimensional, three-dimensional. |  |
| dsDNA | Double stranded DNA. |  |
| ssDNA | Single stranded DNA. |  |
| c-ssDNA | Complementary single stranded DNA, which can form dsDNA. |  |
| cc, icc, tcc | Correct-contact, incorrect contact and trap correct contacts. |  |
| $l_p$ | $= 3.4 \times 10^{-10} \text{ m} = 1 \text{ bp}$, Distance between two consecutive bases of dsDNA. | 1 bp |
| $L, u$ | Length of the template and probe ssDNA polymers. | Nucleotides, nt |
| $v$ | Volume of a nucleotide monomer. | $\text{bp}^3$ |
| $V$ | Reaction volume which is equal to the sphere with radius equal to the sum of the radii of gyration of c-ssDNAs. | $\text{m}^3$ |
| $vL$ | Volume of the ssDNA polymer of length $L$. | $\text{m}^3$ |
| $c$ | Complexity of DNA sequence. Number of unique bases present in the given DNA sequence. | nt |
| $\langle n_{cc} \rangle$ | $(= vL / V)$, average number of correct contacts between the complementary ssDNA strands when they are confined within a reaction volume $V$. | dimensionless |
| $\langle n_{icc} \rangle$ | $(= vL^2 / V)$, average number of incorrect contacts formed between the complementary ssDNA strands when they are confined within a reaction volume $V$. | dimensionless |
| $\langle n_{tcc} \rangle$ | $(= vL^2 / c V)$, average number of trap correct contacts formed between the complementary ssDNA strands with complexity $c$ when they are confined within a reaction volume $V$. | dimensionless |
| $k_t$ | $= k_{sm} \phi$, where $k_{sm}$ is the maximum possible Smoluchowski rate limit and $\phi$ is the factor associated with the electrostatic repulsions at the DNA-DNA interface. | (mol/lit/s) |
| $k_{cc}$ | $= k_t \langle n_{cc} \rangle$, is the number of correct contacts formed upon interpenetration of c-ssDNAs. | (mol/lit/s) |
| $k_{icc}$ | $= k_t \langle n_{icc} \rangle$, is the number of incorrect contacts formed upon interpenetration of c-ssDNAs. | (mol/lit/s) |
| $k_{tcc}$ | $= k_t \langle n_{tcc} \rangle$, is the number of trap correct contacts formed upon interpenetration of repetitive c-ssDNAs. | (mol/lit/s) |
| $k_N$ | Nucleation rate constant that directly proportional to $k_{cc}$. | (mol/lit/s) |
| $k_Z$ | Zippering rate constant. | 1/s |
| $M_1, M_2$ | Concentrations of c-ssDNA strand 1 and 2. | (mol/lit) |
| $M_D$ | Concentration of duplex form DNA. | (mol/lit) |
| $W$ | Concentration of nucleus DNA. | (mol/lit) |
| $Y$ | Concentration of incorrect contacts. | (mol/lit) |
| $N$ | Size of the nucleus. | nt |
| $R_g$ | Average radius of gyration of c-ssDNA or SARW. | bp |
| $q, q_t$ | Slithering lengths associated with the complementary ssDNAs with icc and tcc. | nt |
| $k_r, k_{rt}$ | Dissociation rates associated with c-ssDNAs having icc and tcc respectively. | 1/second |
| $k_R$ | Overall renaturation rate. $k_{RQ}$ where Q = I, II, III and IV correspond to the respective renaturation Schemes I, II, III and IV. | (mol/lit/s) |
| $k_+, k_-$ | Microscopic forward and reverse zipping rate constants. | nt / second |

Here *v* / V is the probability of finding any nucleotide of c-ssDNAs inside *V* upon each movement and *vL* is the total chain volume. Upon assuming c-ssDNAs as a spherical shaped nucleotide clusters, the reaction space can be thought approximately as a sphere with radius of 2*R_g_* where *R_g_* is the average radius of gyration of the individual c-ssDNA. One can straightforwardly interpret these results as follows. Since *L* number of monomers of the c-ssDNA of interest search for the complementary nucleotides on the other strand at the same time, one obtains 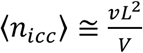. Out of these *n_icc_* numbers of incorrect contacts, the probability of finding the trap correct contacts among the repetitive c-ssDNAs will be 1 / c. Therefore, one obtains 〈*n_tcc_*〉 = 〈*n_icc_*〉/*c*. In the same way, the probability of finding the correct contact will be 1 / *L* that is independent on the number of repeats. From this one can deduce that 〈*n_cc_*〉 = 〈*n_icc_*〉/*L*. This expression for 〈*n_tcc_*〉 will work only when *L* is much higher than the sequence complexity *c* so that 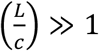. When *c* → *L*, then there are always chances for the occurrence of partial repeats and one can obtain the approximation 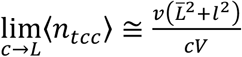 where 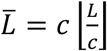 is the length of c-ssDNAs containing full repeats and 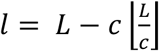 is the length of c-ssDNAs with partial repeat. For example, when *L* = 20 and *c* = 6, then one obtains 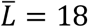 and *l* = 2. Here 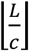 is the floor function operator. For example, 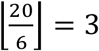. When the main template strand length is fixed at *L* and only the length of the probe *u* is varied as in case of PCR reactions, then we find the following scaling relationships.

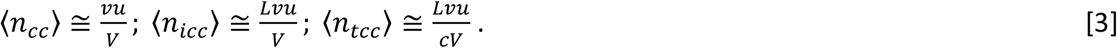

When the length of both c-ssDNAs are randomized within (0, *L*) with equal probabilities (= 1 / *L*) resembling the uniformly sheared DNA [8] so that the average length will be <*L*> = *L* / 2, then one finds the following scaling relationships.

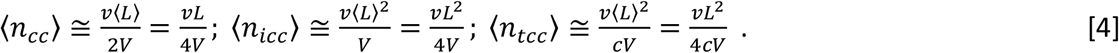

When the complexity *c* = *L*, then irrespective of the sheared nature of c-ssDNAs, the probability of finding the correct contact will be *p_cc_* = 1 / *L*. As a result, one obtains 〈*n_cc_*〉 ≅ *p_cc_*(*n_icc_*〉 = *Lv*/4*V* for the case of sheared DNA. When *c* → 〈*L*〉 then one can obtain the approximation 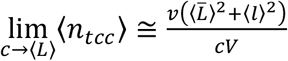 where 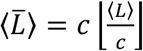 and 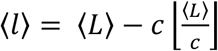 by definition as in case of non-sheared and repetitive c-ssDNAs.

### 2.3. Radius of gyration of c-ssDNAs

The ensemble average of the radius of gyration of a chain molecule can be defined as 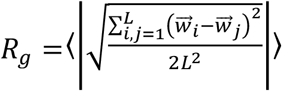 where 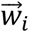 is the vector coordinates (*X, Y, Z*) of the i^th^ monomer and *L* is the length [38,39]. Here the averaging is done over several possible spatial configurations of the polymer. Detailed numerical simulations suggested that the scaling of *R_g_* with *L* deviates significantly from the standard scaling 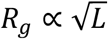 when the ratio 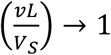 where *V_S_* is the volume in which the SARW is confined. In other words, the scaling 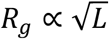 will be valid only in the limit 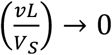. In this context, nonlinear least square fittings of the values of *R_g_* obtained over various lengths of SARWs confined in different volumes *V_S_* using Marquart-Levenberg algorithm [40] suggested the following functional form.

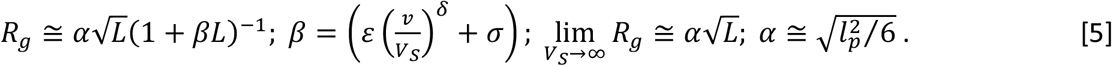

Here α is the pre-exponent and *l_p_* is the physical distance between the monomers. Clearly, in the limit as *V_S_* → ∞ that represents the dilute *in vitro* conditions, we recover the well-known scaling relationship for a linear chain polymers as 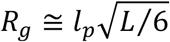. The excluded volume effects will be prominent when the intrinsic volume of the polymer *vL* is close to the confinement volume *V_S_* [8]. Under such conditions, the interpenetration of c-ssDNAs among each other will be very much limited. In our model, c-ssDNAs are not physically confined in space. Rather, they are allowed to enter or exit the reaction volume freely. Hence, we assume 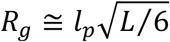 and subsequently derive the following expressions for the reaction and monomer volumes.

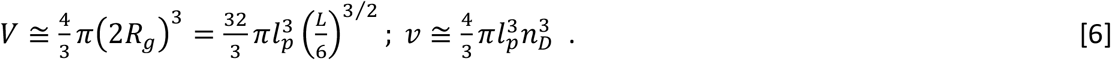

In these equations, *l_p_n_D_* is the radius of gyration of the nucleotide monomer in terms of number (*n_D_*) of distances between adjacent nucleotides (*l_p_* ≅ 3.4 × 10^−10^ m for dsDNA and *l_p_* ≅ (5.9 to 7) × 10^−10^m for c-ssDNAs, since *l_p_* corresponding to c-ssDNAs is a fluctuating quantity, we denote *l_p_* corresponding to dsDNA as the standard base-pair unit, bp) and the volumes are measured in bp^3^. When we model DNA as chain of beads where each bead represents a monomer, then the radius of the monomer bead will be approximately equal to the radius of the DNA cylinder which is approximately equals to *n_D_l_p_* ≈ 3 bp. Using **Eqs. 6**, when *L* ≫ *c*, then one can derive the following expressions for the number of cc, icc and tcc formed upon interpenetration of c-ssDNAs inside the reaction volume.

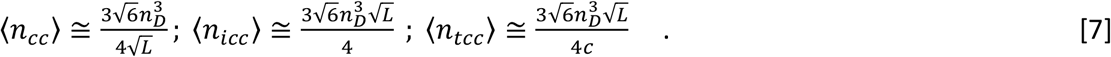

When c-ssDNAs are equal in length, then using **Eqs. 6** and **7** one can drive expressions for the overall rate associated with the formation of cc, icc and tcc as follows.

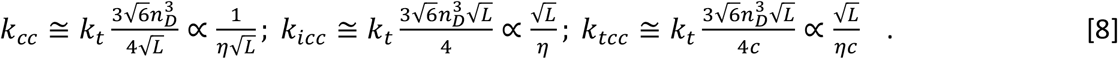

### 2.4. One step DNA hybridization model

Using the scaling results presented in **Eqs. 7** and **8**, we now revisit the Wetmur-Davidson model [8]. When the nucleation occurs via pure 3D diffusion controlled collision route as described in **Scheme I** of **Fig. 1A**, then the nucleation occurs with a rate *k_N_* ∝ *k_cc_* and one can straightforwardly derive the scaling as 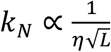. This means that, *k_N_* = *ζk_cc_* where *ζ* is the dimensionless proportionality constant. In this background, the differential rate equations for the renaturation of a nonrepetitive c-ssDNAs can be written as follows.

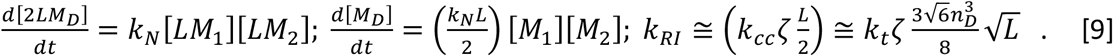

In this equation, we have substituted *k_N_* = *ζk_cc_*. Here *k_RI_* is the overall hybridization rate corresponding to Scheme I, M_D_ (mol/lit) is the concentration of dsDNA molecules and *M_1_* and *M_2_* (mol/lit) are the concentrations of the c-ssDNA molecules. Upon multiplying by *L* (or 2L in case of dsDNA) one can convert these concentrations of the polymer molecules into the concentrations of nucleotides in each category which are actually the experimentally observed variables. In the derivation of **Eqs. 9**, one assumes that the zipping is spontaneous and the nucleation is the rate limiting step. Remarkably, **Eqs. 9** associated with the Wetmur-Davidson model correctly predicts the length dependent scaling as 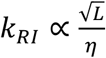 in line with the experimental observations [8,14].

### 2.5. Two step mechanism of DNA hybridization

When the concentration of the nucleus is *W* and the average size of a nucleus is *N* nt (so that the concentration of nt in the nucleus form dsDNA is [2*NW*]), then from the two-step mechanism given in **Scheme II** of **Fig. 1A**, one can derive the following rate equations.

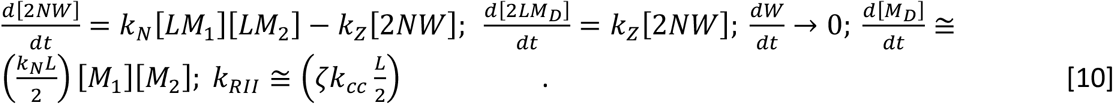

In this equation, *k_RIII_* is the overall hybridization rate corresponding to Scheme II, *k_Z_* (nt/s) is the zipping rate constant. When zipping is much faster than the nucleation step, then the rate equations given in **Eqs. 10** attain steady state so that one can set 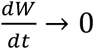. Here nucleation will be the rate limiting step. In such scenario, **Eqs. 10** reduce to **Eqns. 9** where *k_cc_* = *k_icc_/L* for nonrepetitive c-ssDNA. Therefore one obtains the expression 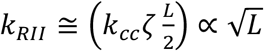.

### 2.6. Limitations of one step and two-step hybridization models

The main flaws of **Eqs. 9** and **10** are as follows.

1. Clearly, **Eqns. 9** and **10** will work only for short DNA segments for which one can ignore the zipping times. However, for long c-ssDNAs, the zipping time τ_Z_ scales with the length of DNA in a linear or square [15] or power law manner [27]. In general, one observes the scaling for the zipping rate (which is the inverse of the zipping time) as *k_Z_* ∝ *L*^−*ρ*^. There are two extreme possibilities. **1**) When the zipping is diffusion like and not energetically driven or hampered by the chain entropy of the single stranded overhangs, then the zipping will be a diffusion like process and *k_Z_* ∝ *L*^−2^. **2**) When the zipping is irreversible and energetically driven process over the entropic barriers, then one finds that *k_Z_* ∝ *L*^−1^. In reality, the chain entropy barrier decreases in the process of zipping and the stability of duplex increases along the transition from ssDNA to dsDNA form. In general, when the entropic barrier associated with the single strand overhangs is significant then one observes the exponent as 1≤ *ρ* ≤ 2 depending on the type of prevailing conditions. For example, when the zipping is similar to that of the forced translocation of a polymer through a nanopore [27], then one finds that *ρ* ≅ 1.37.
2. When *L* = 1, then one finds from the extrapolation of experimental data [21] that 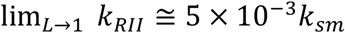 where *k_sm_* = *k_t_* / *ϕ* is the maximum possible Smoluchowski diffusion controlled bimolecular collision rate limit across neutral molecules. However, **Eqs. 9** and **10** associated with the 3D diffusion only model predict that 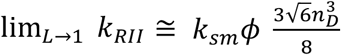. This means that the contributions from the electrostatic repulsions across the negatively charged phosphate backbones over the rate enhancement at the DNA-DNA interface of c-ssDNAs should be 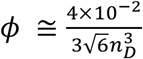 where *n_D_* ≅ 3 nt.
3. When the mode of nucleation is via pure 3D diffusion, then the number of correct-contacts or the nucleation rate should be independent of the number of repeats in the c-ssDNA strands. Since the zipping step directly follows from the 3D diffusion mediated nucleation, the overall renaturation rate should be independent of the complexity of c-ssDNAs. Further, the presence of trap correct-contacts would slow down the nucleation process since tcc across the c-ssDNAs need to be broken before exploring other locations for the correct-contacts [13,25]. This incurs significant amount of time lapse which in turn delays the renaturation process. On the other hand, such tcc keeps the complementary strands in the close vicinity for an extended amount of time compared to icc. This is essential for the efficient operation of various one dimensional facilitating processes such as slithering, inchworm movements and internal displacement mechanisms [13,25] which can speed up the rate of nucleation. Detailed molecular dynamics simulations also reveals that the metastable states arising out of trap correct contacts among the short repetitive c-ssDNAs can significantly speed up the hybridization process [33].
4. For repetitive ssDNA with complexity *c* and *L* / *c* number of repeats, Wetmur-Davidson model directly assumed the scaling 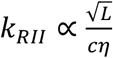 which predicted that 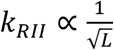 at *c* = *L* that is not consistent with their experimental observations [8,14]. This critical flaw and other arguments arising out of the simulation results [25] clearly suggest that the nucleation process must involve at least two different pathways viz. 3D diffusion only mode and a mode with the combination of 1D and 3D diffusions [15] as described in **Schemes III** and **IV** of **Fig. 2**. Here 3D diffusion only pathway progresses directly via correct-contacts and it works only for short c-ssDNAs where the zipping times can be ignored. The 3D-1D diffusion pathway can progress via either incorrect contacts or trap correct contacts depending on the repetitive nature of c-ssDNAs.

### 2.7. Three step models of DNA hybridization

According to **Scheme III**, the base clusters of c-ssDNAs interpenetrate each other upon collision to the form incorrect contacts in the first step with a rate k_icc_. Subsequently, the correct contact will be formed via various 1D facilitating processes such as slithering, inchworm movements and internal displacement mechanisms. These are all analogous to the facilitating processes such as sliding, hopping and intersegmental transfers exhibited by the DNA binding proteins in the process of searching for their cognate sites on DNA [41]. Nucleation occurs upon finding the correct-contacts over several rounds of incorrect-contact formation, 1D slithering movements and dissociations. Remarkably, detailed molecular dynamics simulation on the hybridization of short ssDNA segments also revealed the presence of four state (three-step) mechanism [33]. With this background, **Eqs. 10** can be rewritten for the case of nonrepetitive c-ssDNAs as follows (Scheme III in **Fig. 2**).

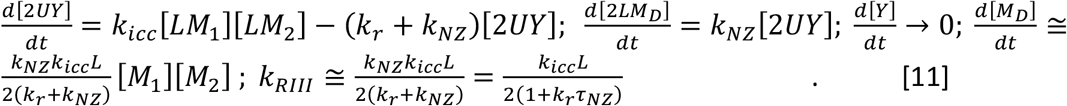

In these equations, *k_RIII_* is the overall hybridization rate corresponding to Scheme III, [2*UY*] is concentration of the nucleotides involved in the icc, *k_r_* (1/s) is the dissociation rate constant connected with the icc, 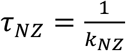 is the total time required for the nucleation and zipping through incorrect-contact formation route and its inverse will be the overall nucleation-zipping rate. After each incorrect-contact, c-ssDNAs perform 1D slithering on each other over *q* nucleotides on an average and then dissociate. The time required for such 1D diffusion will be 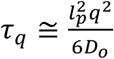 [15]. In this time, c-ssDNAs scan *q* nt on each other for the presence of correct contacts. To completely scan the entire c-ssDNAs of length *L*, at least *L* / *q* numbers of such cycles of incorrect-contact formation, 1D slithering and dissociations are required. Therefore, the nucleation time can be expressed as 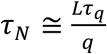 which is the minimum amount of time that is required by the c-ssDNAs to find a correct-contact. When c-ssDNAs are nonrepetitive, then one finds the total time required for the nucleation and zipping steps as *τ_NZ_* ≅ *τ_Z_* + *τ_N_* where 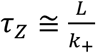 and 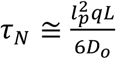 are the zipping and nucleation times through the incorrect contact route [15]. The nucleation rate will be the inverse 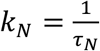 and the zipping rate will be 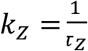. Noting that fact that nucleation will be immediately followed by zipping, we combine the nucleation and zipping times.

Since the stabilizing effects of the already formed Watson-Crick base pairs are much stronger than the entropic barriers arising out of the freely moving single stranded overhangs, one can assume the linear scaling for the zipping time with the length of c-ssDNAs as *τ_Z_* ∝ *L*. Here *k*_+_(nt/s) is the microscopic zipping rate (**Fig. 2**), *l_p_* = 1 bp, *q* (nt) is the average slithering length and *D_O_* (bp^2^/s) is the phenomenological 1D diffusion coefficient associated with the slithering dynamics of c-ssDNA segments. Upon inserting the expressions for the nucleation and zipping times into the expressions for *k_RIII_* as given in **Eq. 11** one obtains the following result.

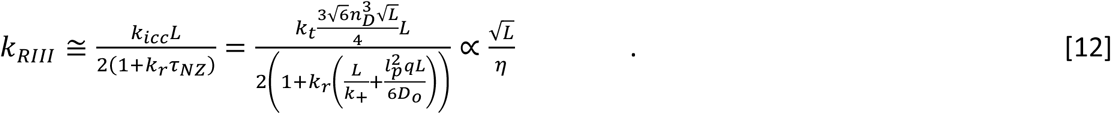

Here one should note that *D_O_* will be influenced only by the local viscosity at the DNA-DNA interface of the c-ssDNAs with icc or tcc and it will not be much influenced by the global viscosity that is connected with the translational diffusion of the entire c-ssDNA nucleotide cluster. In the presence of repetitive sequences, apart from icc and cc, trap correct contacts are also formed as shown in **Scheme IV** of **Fig. 2**. Clearly, the nucleation can occur via both icc and tcc in case of repetitive c-ssDNAs which is also substantiated by the coarse grained molecular dynamics simulations [29]. Particularly, these tcc can influence the renaturation in two possible ways viz. **1**) they keep the complementary strands in close vicinity for prolonged amount of time that enhances the slithering times which in turn increases the efficiency of searching for the correct contacts [29,33]. **2**) these kinetic traps prolong the overall renaturation timescales which will be more prominent especially when the length of ssDNA is very long. As a result of these, the dissociation rate connected with the partial duplexes decreases (*k_r_* → *k_rt_*) and the slithering length increases (*q* → *q_t_*). The overall renaturation rate corresponding to the tcc route can be obtained by replacing (*k_r_* → *k_rt_*, *q* → *q_t_*, *k_icc_* → *k_tcc_*) in **Eq. 12**. Upon combining the contributions of these parallel pathways through icc and tcc routes, one finally obtains the following expression for the overall renaturation rate for the repetitive c-ssDNAs.

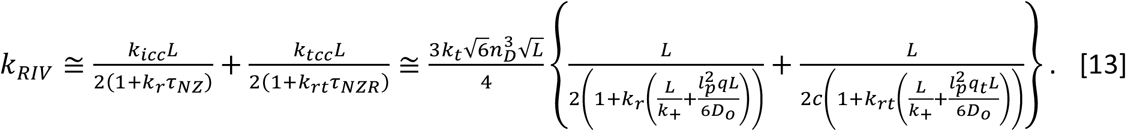

**Eq. 13** is the central result of this paper. In this equation, *k_RIV_* is the overall hybridization rate corresponding to the repetitive c-ssDNAs as described by **Scheme IV**, *τ_NZR_* ≅ *τ_Z_* + *τ_NR_* is the nucleation-zipping time via trap correct contact route where 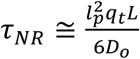 is the average nucleation time, *k_rt_* is the dissociation rate connected with tcc and *q_t_* is the 1D slithering length associated with the repetitive c-ssDNAs with partial duplexes and single strand overhangs. The nucleation rate will be the inverse 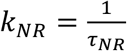. When *L* > *c*, then *k_r_* > *k_rt_* since the partial duplexes are more stable than the incorrect contacts and *q* < *q_t_* since the c-ssDNAs stay close to each other for prolonged amount of time which allows extended amount of slithering. As a result these, we find that 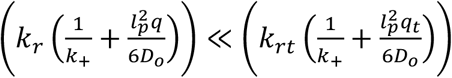 from which one obtains the generally observed scaling as 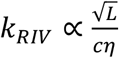. Remarkably, when *c* = *L*, **Eqs. 13** predicts the correct scaling 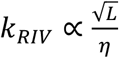 in line with the experimental observations [8,14].

**Eq. 13** will be still valid when c-ssDNAs are unequal in length similar to that of the template and probe DNAs used in the PCR reactions. However, in this case the length of duplex formed upon hybridization will be equal to the length of the probe. Let us denote the lengths of template and probe as *L* and *u* respectively. When *L* ≫ *u*, then the template strand diffuses slower than the probe strand and one can show that the bimolecular collision rate associated with the template-probe system scales as 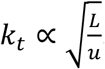. This follows from the fact that 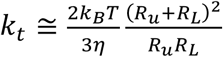 where 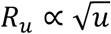 and 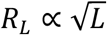 are the radius of gyration of the probe and template strands respectively [35]. When *u* ≪ *L*, then the reaction volume will be almost independent on the volume of the probe. This results in the scaling as 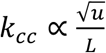 for the rate of formation of the correct contacts. Since the nucleation rate is directly proportional to the rate of formation of the correct-contacts across c-ssDNAs, this means that the nucleation rate scales with the lengths of the probe and template c-ssDNAs as 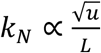 since *k_N_* ∝ *k_cc_* which reduces to the observed scaling corresponding to the nucleation rate 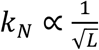 as *c* → *L*.

## 3. Simulation Methods

To understand various scaling relationships associated with the correct, incorrect and trap correct contacts in the process of renaturation, we modelled the c-ssDNAs as self-avoiding random walks confined inside a 3D lattice box that mimics the reaction volume. When the box is a cube with side *b*, then the box volume will be *V* = *b*^3^ cubic units and (*X, Y, Z*) = (0, *b*) are the reflecting walls for the generation of SARW 1 and 2 which mimic c-ssDNA strands 1 and 2. The SARWs were generated starting from a random point by sequentially calling three random number *r*_1,2,3_ ∈ (0,1). If *r*_1_ < 0.5, then X→*X* + 1 else *X* → *X*– 1 and similar rules were set for *Y* and *Z* dimensions. The move will be allowed only when the next location is not self-intersecting the earlier locations which mimics the implementation of the excluded volume effects. SARW 1 and SARW 2 cannot self-intersect but they can cross-intersect each other that mimics the interpenetration and contact formation among the pair of c-ssDNAs.

When the sequence is repetitive with complexity *c*, then in such a polymer of length *L* > *c*, the positions from 1 to *c*, *c* + 1 to 2*c* and 2*c* + 1 to 3*c* and so on are identical in sequence. Those contacts between c-ssDNAs without exact registry matches are the incorrect contacts. The positions from 1 to *c* of SARW 1 can form duplex with position stretch *c* + 1 to 2*c* of SARW 2 and so on. These cross intersections can be classified as correct contact (cc), incorrect contact (icc) or trap correct-contact (tcc) depending the position of the registry match. In this setting, the volume of a SARW with length *L* will be *vL* where *v* = 1 is the volume of the monomer unit. In a repetitive c-ssDNA with complexity *c*, when there is a contact between position 1 of SARW 1 with position c + 1 of SARW 2, then it is defined as a trap correct contact. When the cross-intersection points are exact registry matches, then they are the correct-contacts. For example, when there is a contact between position 2 of SARW 1 with position 2 of SARW 2, then it is a correct contact. All other types of contacts are incorrect-contacts by definition.

To understand the effects of sheared c-ssDNAs on the overall rate of renaturation, we randomized the lengths of SARWs. Length of the original c-ssDNA is L and we denote the index of the monomers from 1 to L. Shearing of c-ssDNAs will generate fragments of this template strand with random lengths. To mimic this, two random integers r_1_, r_2_, were generated inside (1, L) which are the random lengths of SARW 1 and SARW 2. The starting point of SARW 1 was defined by calling a random integer si inside (1, L-r_1_) and the end point will be s_1_ + r_1_. Similarly the starting point of SARW 2 was defined by calling random integer s_2_ inside (1, L-r_2_) and the end point will be s_2_ + r2. These start and end locations of SARW 1 and 2 were used to compute the number of correct, incorrect and trap correct contacts.

Here we assume that the pair of c-ssDNAs have already reached the reaction volume *V* via 3D translational diffusion with a bimolecular collision rate *k_t_*. The quantities that we are interested to compute here are the average number of correct, incorrect and trap correct contacts formed between a pair of c-ssDNAs of length *L* and complexity *c* inside the reaction volume *V*. To achieve this, several pairs of SARWs were generated inside a closed cubic lattice box as in **Fig. 3** and the number of cc, icc and tcc were enumerated and averaged over 10^5^ trajectories. The average number of cc, icc and tcc seem to be influenced by the length of c-ssDNA (*L*), its sequence complexity (*c*) and the reaction volume (*V*). To investigate the effects of these variables, we iterated one factor over a range of values at a time by fixing other two factors at constant values. In the real situations, one can translate the results of the lattice model by setting the reaction volume as the volume of the sphere with radius equals to 2*R_g_* where *R_g_* is the average radius of gyration of c-ssDNAs. To understand the distribution of icc, cc and tcc, we constructed the histogram of samples drawn from large population of counts over 10^4^ numbers of SARW trajectories.

**FIGURE 3.**
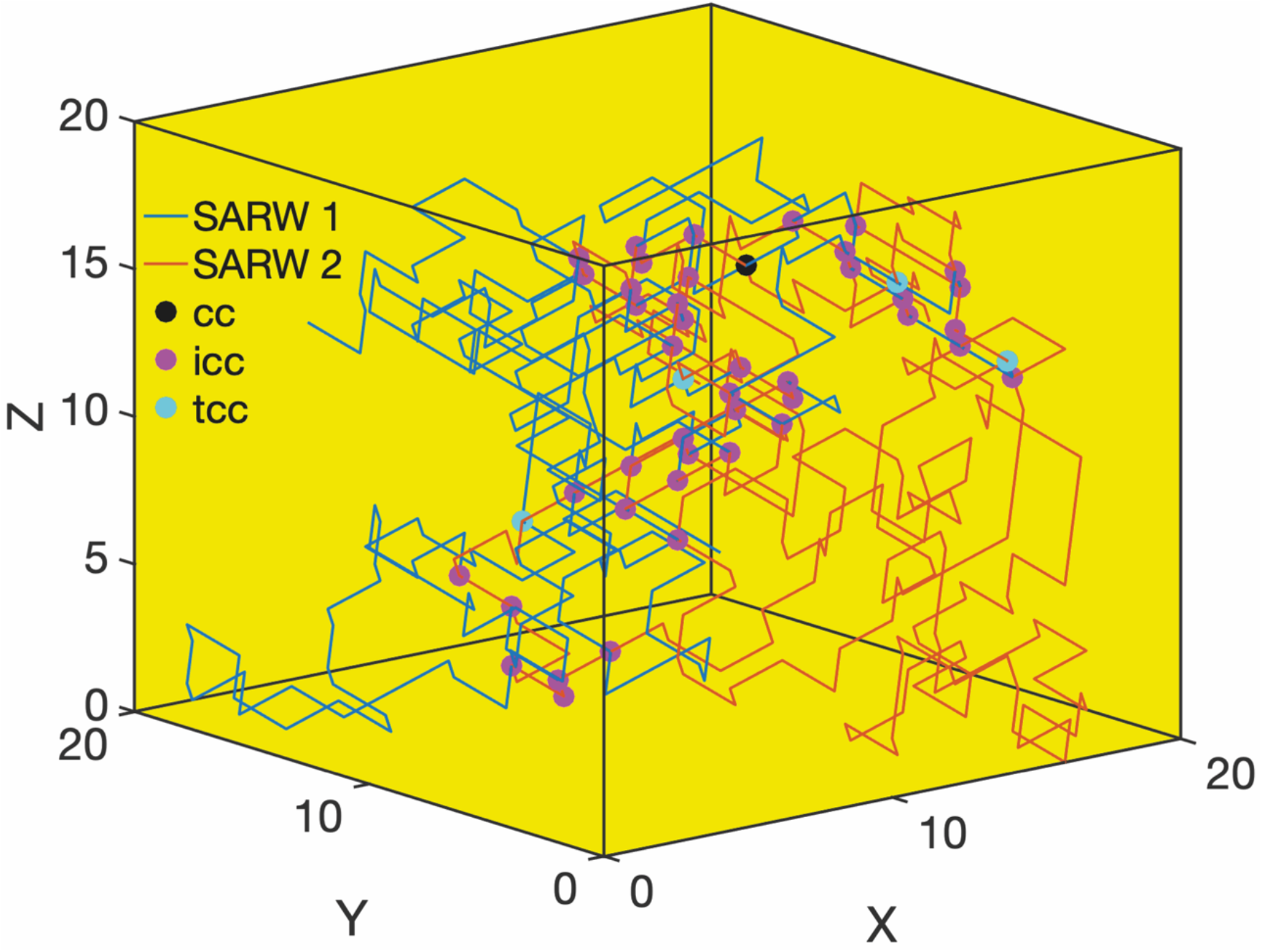
Lattice model on DNA renaturation. Here the complementary single strands are modelled as self-avoiding random walks (SARW 1 (blue) and 2 (red)) confined inside the reaction volume defined by the lattice box. The position of the nucleotides are marked from 1 to *L*. Both the strands arrive at the reaction volume via 3D diffusion. Correct contact between these SARWs occurs when there is an exact registry match that can further leads to nucleation and zipping. For example, when there is a contact between position 7 of SARW 1 with position 7 of SARW 2, then it is defined as a correct-contact (cc). When there is a contact between position 7 of SARW 1 with position 5 of SARW, then it is an incorrect contact (icc). When the complexity is *c* < *L*, then the SARW has *L* / c number of repeats i.e. the sequence spanned across position 1 to *c*, c + 1 to 2c and 2*c* + 1 to 3*c* and so on will be the same. Therefore, when there is a contact between position 1 of SARW 1 with position *c* + 1 (or 2c + 1 and so on) of SARW 2, then it is defined as a trap correct-contact (tcc). Here tcc can lead to the formation of partial duplexes with single stranded overhangs which are actually kinetic traps in the renaturation pathway. Here the settings are, *L* = 250, *c* = 5, and reaction volume *V* = 20^3^, number of incorrect contacts (*n_icc_*) = 25, trap correct contacts *n_tcc_* = 4 and the number of correct contacts *n_cc_* = 1 that occurs at (*X*, *Y, Z*) = (9, 8, 17).

## 4. Results and Discussion

The c-ssDNA strands can be modelled as self-avoiding random walks with the average radii of gyration *R_g_*. Renaturation of c-ssDNAs requires the formation of correct contacts between them that leads to nucleation and zipping. Here c-ssDNAs reach the reaction volume via 3D translational diffusion. Upon entering the reaction volume, they can interpenetrate each other to form various types of contacts such as cc, icc, tcc or no contact at all. Clearly, when the size of c-ssDNAs are approximately equal then the rate at which they enter the reaction volume via translational diffusion 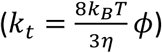 will be independent of the radius of gyration. However, the extent of interpenetration of these strands will be strongly dependent on the length of c-ssDNAs *L* and the confinement volume *V*.

Detailed lattice model simulations revealed the general scaling of various types of contacts between c-ssDNA strands as 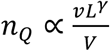 for nonrepetitive c-ssDNAs where Q = (icc, cc) and *v* is the monomer volume. For cc, we find the exponent *γ* = 1 and for icc we find that *γ* = 2. For repetitive c-ssDNAs one finds that 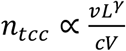 with *γ*= 2. When *c* = *L*, this reduces to 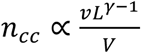 resembling the nonrepetitive case. Variations of the number of tcc, cc and icc with respect to changes in *L* with fixed *V* and *c* are demonstrated in **Fig. 4**. Variations of the number of tcc, cc and icc with respect to changes in *V* with fixed *L* and *c* are demonstrated in **Fig. 5**. Results clearly suggest that the scaling 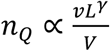 where Q = (icc, cc) is independent on the shape and dimension of the confining lattice box as shown in **Figs. 4B-D** and **5B-D**. Similar results for the repetitive c-ssDNAs are demonstrated in **Fig. 6A**. We can conclude from these results that the number of correct contacts scales with *L* and *V* as 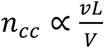 irrespective of the complexity of c-ssDNA strands and shape and dimension of the confining lattice box.

**FIGURE 4.**
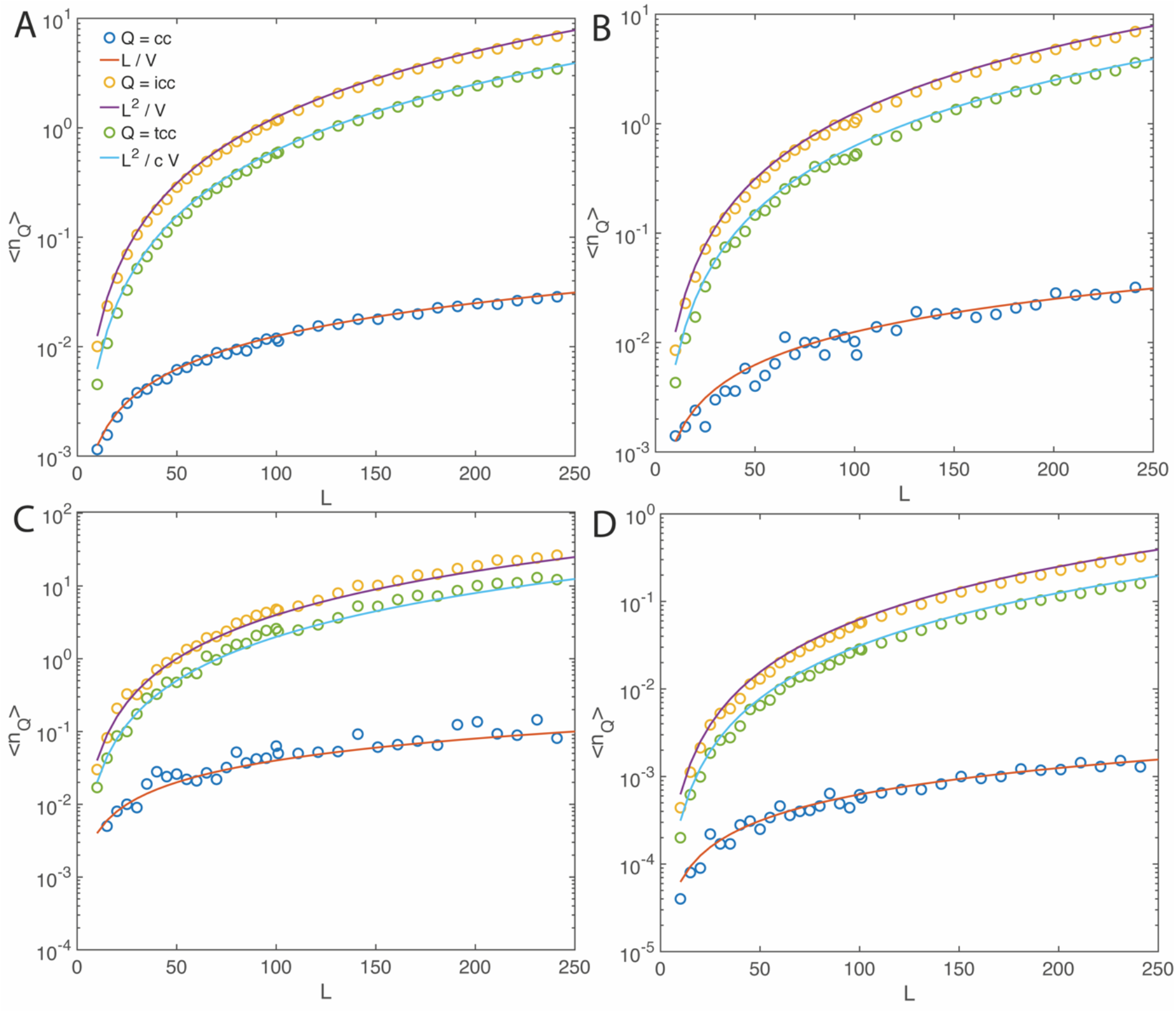
Variation of the average number of correct (*n*_cc_), incorrect (*n_icc_*) and trap correct contacts (*n_tcc_*) with respect to the length L of SARWs and dimensionality. Average number of various contacts were obtained over 10^5^ SARW trajectories. Hollow circles are the stochastic simulations results and solid lines are the predictions by **Eqs. 2**. Irrespective of the dimensionality and shape of the lattice box where the volume of the monomer unit is *v* = 1, the expression given in **Eqs. 2** are valid. **A**. Three dimensional SARW confined in volume *V* = 20^3^, length was iterated in L = (10, 250) and *c* = 2. **B**. Three dimensional SARW confined inside a box with base area = 10^2^ (10 by 10) and length = 80 so that *V* = 8000. The length of SARW is iterated inside *L* = (10, 250) with complexity *c* = 2. **C**. two dimensional SARW confined in *V* = 50^2^, and the length of SARW was iterated inside *L* = (10, 250) and *c* = 2. **D**. Four dimensional SARW confined in *V* = 20^4^, *L* = (10, 250) and *c* = 2.

**FIGURE 5.**
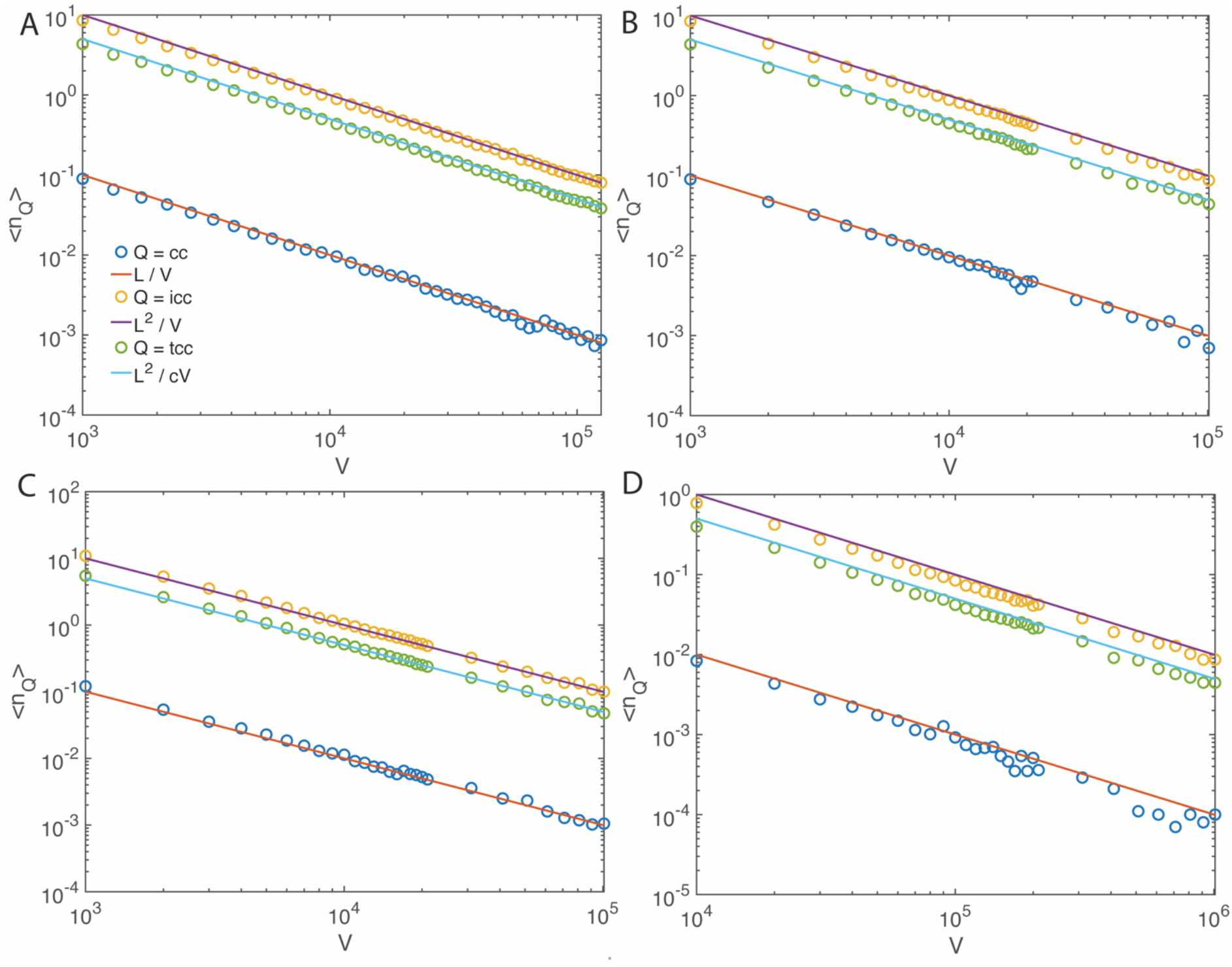
Variation of the average number of correct (*n_cc_*), incorrect (*n_icc_*) and trap correct contacts (*n_tcc_*) with respect to change in confinement volume of SARW and dimensionality. Average number of various types of contacts were obtained over 10^5^ trajectories. Hollow circles are the stochastic simulations results and solid lines are the predictions by **Eqs. 2**. Irrespective of the dimensionality and shape of the lattice box where the volume of the monomer unit is *v* = 1, the expressions given in **Eqs. 2** are valid. **A**. three dimensional SARW confined in cubic box with volumes iterated inside *V* = (10^3^, 50^3^), length of SARW was set as *L* = 100 and *c* = 2. **B**. three dimensional SARW confined inside a box with base area = 10^2^ and length iterated from 10 to 1000 so that *V* varies from 10^3^ to 10^5^. The length of SARW is *L* = 100 with complexity *c* = 2. **C**. two dimensional SARW confined box with base side = 100 and length iterated from 10 to 1000 so that *V* varies from 10^3^ to 10^5^, *L* = 100 and *c* = 2. **D**. Four dimensional SARW confined in a box with base 10^3^ (10 x 10 x 10) and other side iterated from 10 to 1000 so that the volume *V* varies from 10^4^ to 10^6^, length of SARW was set as *L* = 100 and the complexity was set as *c* = 2.

**FIGURE 6.**
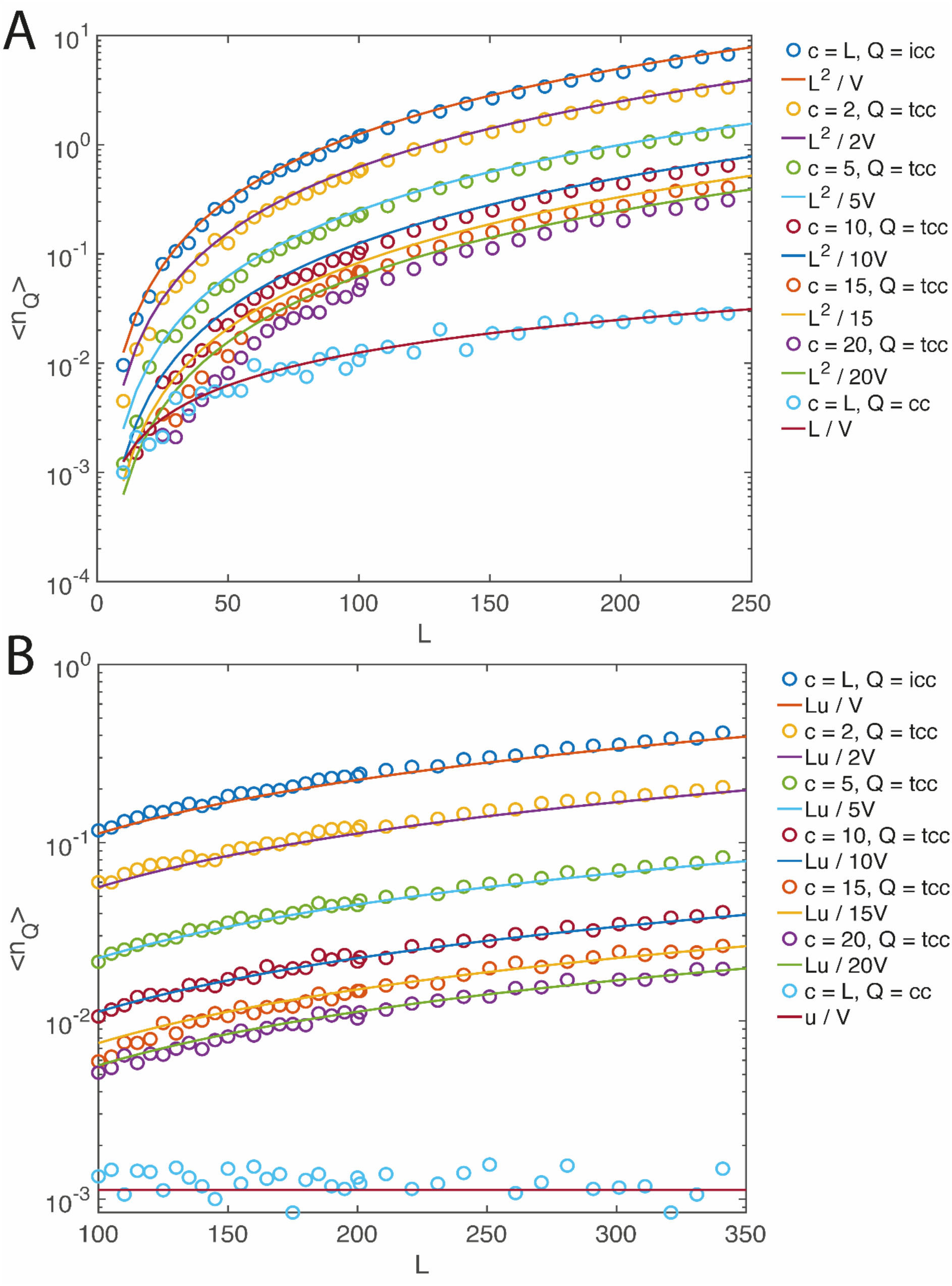
Variation of the average number of correct (*n*_cc_), incorrect (*n_icc_*) and trap correct contacts (*n_tcc_*) with respect to changes in length L and complexity c of SARWs. Average number of various types of contacts were obtained over 10^5^ trajectories Hollow circles are the stochastic simulations results and solid lines are the predictions by **Eqs. 3**. Here the volume of the monomer unit is set as *v* = 1. **A**. the volume is fixed at *V* = 20^3^. **B**. the template strand length is fixed at *L* = 100 and the length of probe is fixed at *u* = 10. The probe was set to form duplex with the template at position stretch 40 to 50. This is the recognition stretch of the probe ssDNA. Both these strands are embedded inside the volume *V* = 20^3^.

These scaling relationships are not affected when the lengths of c-ssDNAs are unequal as demonstrated in **Fig. 6B** or random as demonstrated in **Fig. 7**. Particularly, when the length of the template ssDNA is *L* and probe ssDNA is *u* as in case of annealing phase of polymerase chain reactions, then the one obtains the scaling 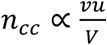. When the c-ssDNA lengths are random with equal distribution inside (0, L), then one observes the scaling 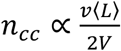 where 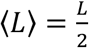 here. These results are demonstrated in **Fig. 7**. Under *in vitro* conditions, *V* will be the reaction volume. Since the reaction radius here is approximately equal to 2*R_g_*, one can consider the reaction volume as 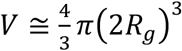.

**FIGURE 7.**
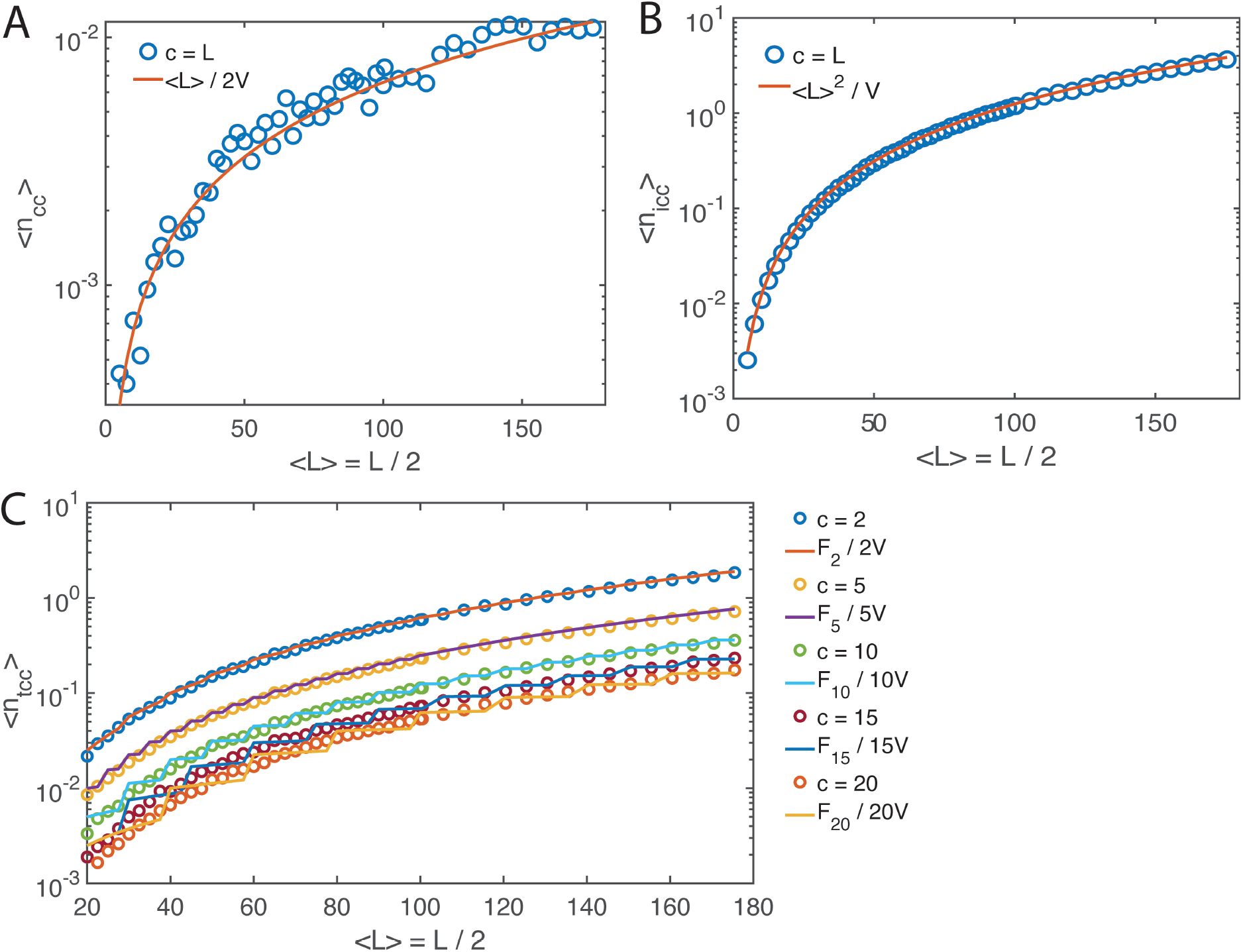
Variation of the average number of correct (*n_cc_*), incorrect (*n_icc_*) and trap correct contacts (*n_tcc_*) with respect to changes in the randomized lengths and complexity *c* of (c-ssDNAs) SARWs. Average number of various types of contacts were obtained over 10^5^ SARW trajectories. Hollow circles are the stochastic simulation results and solid lines are the predictions by **Eqs. 4**. Here the volume is set at *V* = 20^3^ and the length of SARW was chosen randomly within (0, L) with equal probabilities so that the average length is <*L*> = *L* / 2. Here L was iterated inside (10, 360) so that the average length <*L*> will vary from 5 to 180. Results suggest that the probability of finding the correct contact upon collision is 1 / *L* rather than 1 / <*L*> which leads to <*n_cc_*> = *L* / 4*V*. **A**. <n_cc_>. **B**. <n_icc_>. **C**. <n_tcc_>. When the sequence complexity is close to the sequence length, then one finds the approximation 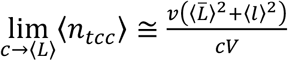 where 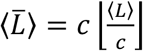 is the length of DNA containing full repeats and 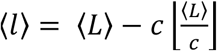 is the length of DNA left over with partial repeat. Here *v* = 1 and 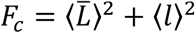. One also should note that **Eqs. 4** will be valid for the repetitive DNA sequences only when (L / c) >> 1.

To understand the effect of varying the confinement volume on the radius of gyration, we iterated the length *L* of SARWs at different confinement volumes. From Flory’s theory, one can conclude that the radius of gyration of the spatially unconfined three-dimensional SARW approximately scales with the length as *R_g_* ∝ *L*^3/5^ [42]. One naturally expects the scaling exponent *θ* in *R_g_* ∝ *L^θ^* as 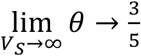. However, when the SARW trajectory is confined inside a cubic box, then the magnitude of the scaling exponent decreases in a complicated manner since the SARW trajectory gets reflected at boundaries of the box which in turn results in tight packaging of SARWs. In this line, several asymptotic functions for the exponent were tried to fit the data on the computed *R_g_* versus the confinement volume of SARWs. However, the volume and length dependency of *R_g_* seems to best fit the functional form 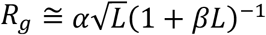 where *β* is a volume dependent parameter. Nonlinear least square fitting using Marquardt-Levenberg algorithm [40] to this function revealed an approximate functional form for the parameter 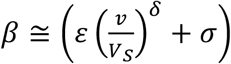 where *v* is the monomer volume, *V_S_* is the confinement volume with the fit parameters *ε* = 0.5 ± 0.04, *δ* = 0.61 ± 0.1 and *σ* = −10^−4^ ± 10^−5^ at 95% confidence level. These results are summarized in **Figs. 8**. The parameter *α* seems to be independent on V_S_ as shown in **Fig. 8B**. Clearly, one obtains the limiting condition 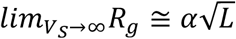 noting the fact that *β* → 0 under such conditions as shown in **Fig. 8C**. Since this limiting condition is valid under *in vitro* conditions, one can use this limiting expression to calculate the molecular and reaction volumes. Remarkably, the numbers of cc, icc and tcc show bimodal type distribution with zero spike as demonstrated in **Figs. 9**. The reason for the zero-spike could be that the confinement volume here is much larger than the intrinsic volume of the c-ssDNA polymer. This mean that the magnitude of the zero-spike is inversely proportional to the volume ratio *vL* / *V*. The number of tcc decreases as the sequence complexity increases which is evident from **Figs. 9C-F**.

**FIGURE 8.**
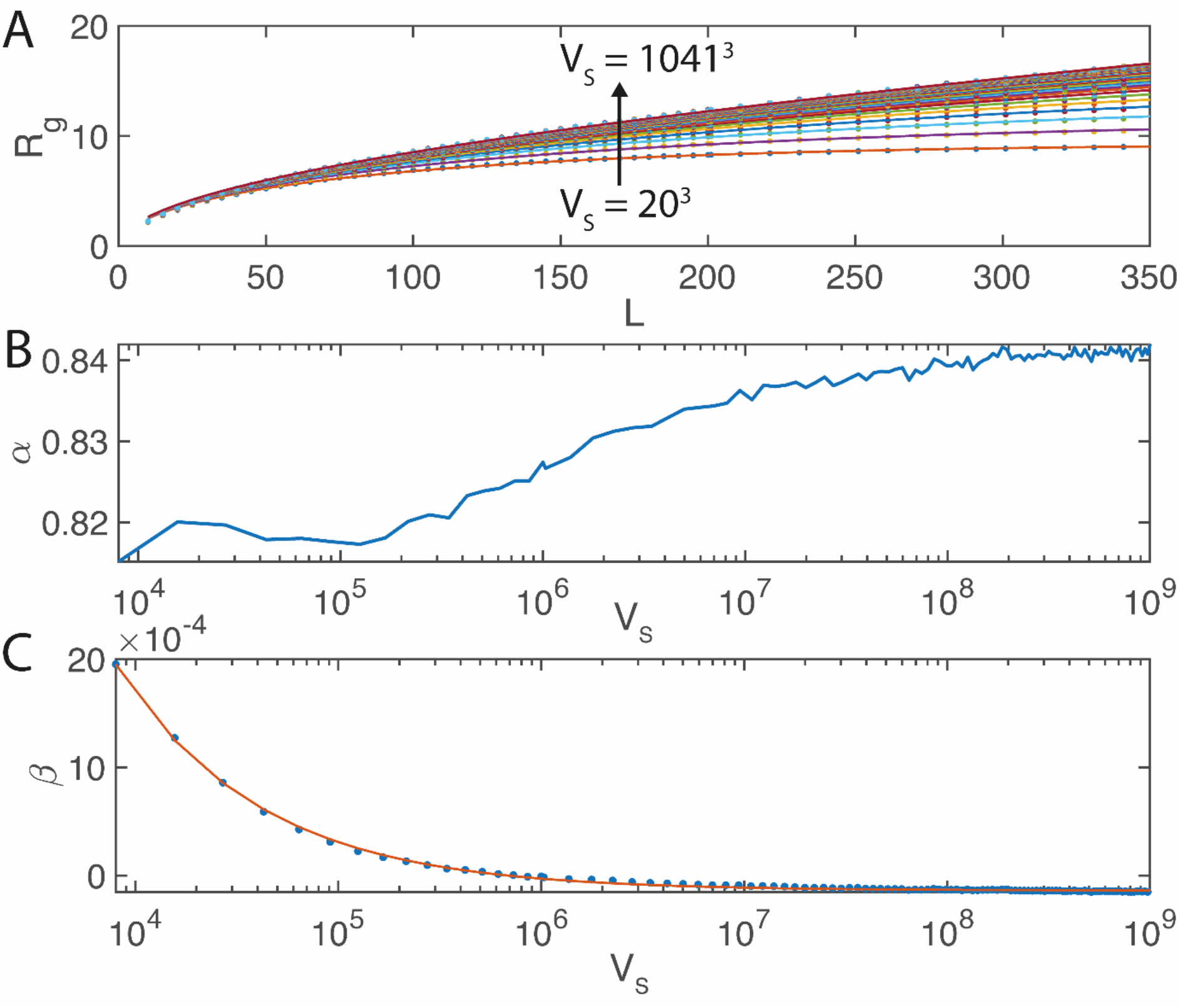
Variation of the average radius of gyration *R_g_* of SARWs with respect to changes in the length *L* and the confinement volume *V_S_* of SARW. The radius of gyration of a chain molecule was calculated as 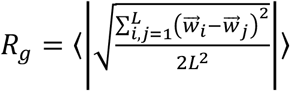 where 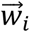 is the vector coordinates (X, Y, Z) of the *i^th^* monomer and *L* is the total length. Average of *R_g_* was obtained over 10^5^ numbers of SARW trajectories. Here filled circles in A and C are the numerical results and solid lines are the nonlinear least square (NLS) fittings. **A**. NLS fits to function 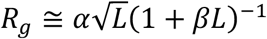 with respect to *L* at various *V_S_* values revealed the parameter *α* that does not change much with *V_S_* as given in **B** and 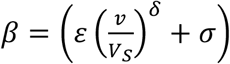 as given in **C** where *ε* = 0.5 ± 0.04, *δ* = 0.61 ± 0.1 and *σ* = −10^−4^ ± 10^−5^. These values were obtained with 95% confidence level. Here the monomer volume is set to *v* = 1 cubic unit.

**FIGURE 9.**
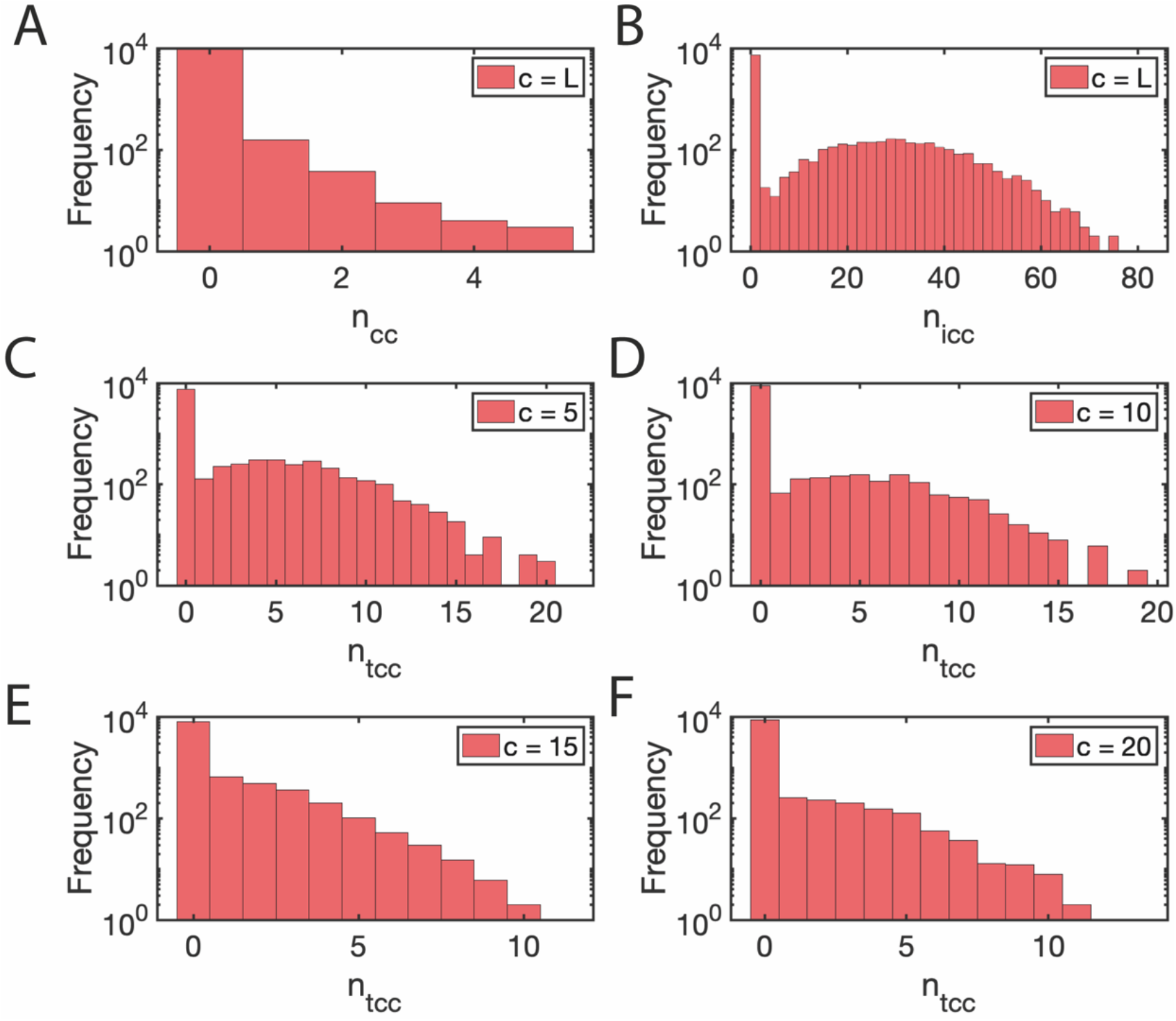
Distributions of the number of correct (*n_cc_*), incorrect (*n_icc_*) and trap correct contacts (*n_tcc_*) at various sequence complexity *c* values. Here volume of the lattice cube is set as *V* = 20^3^, the length of SARW is fixed at *L* = 250 and the sequence complexity c was varied as c = (5, 10, 15, 20, 250). Histograms of various types of contacts were constructed over 10^4^ numbers of SARW trajectories. Clearly, all these distributions show bimodal type with zero spike that represents no contact cases. **A-B**. complexity is set to *c* = 250. **C**. *c* = 5. **D**. *c* = 10. **E**. *c* = 15. **F**. *c* = 20.

When the average radius of gyration of c-ssDNAs scales as 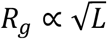, then the reaction volume scales with *L* as *V* ∝ *L*^3/2^. As a result, one observes the scaling associated with the average number of correct contacts formed between the c-ssDNAs upon interpenetration as 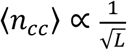. Since the rate of nucleation is directly proportional to the number of correct-contacts, one observes 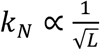 in line with the experimental observations. When c-ssDNAs reach the reaction volume element via 3D translational diffusion, then one finally obtains the viscosity dependence as 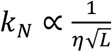. The overall renaturation rate in case of pure 3D diffusion model (Scheme I of **Fig. 1A**) is directly proportional to the nucleation rate as well as *L*. As a result, we finally arrive at the scaling for overall renaturation rate as 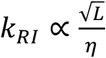 in line with the experimental observations [8] on the hybridization of nonrepetitive c-ssDNAs.

In case of repetitive c-ssDNAs, the average number of correct-contacts is independent on the sequence complexity or the number of repeating elements since it requires only the exact registry match. Clearly, models based on only the 3D diffusion cannot explain the complexity dependence of the renaturation rate. For example, when we assume the scaling 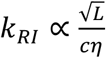 as in case of Wetmur-Davidson model, then we will end up with the inconsistency when *c* → *L* which predicts the scaling 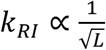 that is not in line with the experimental observations [8]. These arguments clearly suggest that the renaturation follows multiple and parallel pathways in the presence of repetitive sequences. Particularly, the 1D slithering dynamics plays critical roles in the renaturation of the repetitive sequences. The trap correct contacts between the repetitive c-ssDNAs can lead to the formation of partial duplexes with single strand overhangs. Although the entropy component of the single strand overhangs will destabilize these kinetic traps in the pathway of renaturation, tcc keep the c-ssDNAs in the close vicinity for a prolonged timescale that is essential for efficient slithering dynamics and internal displacement mechanisms which in turn accelerates the search for the correct contacts. Clearly, the extent of possible slithering dynamics is directly proportional to the number of trap correct contacts which is inversely proportional to the sequence complexity.

The pathway of renaturation via incorrect contact route always operates irrespective of the presence of repeating elements. Therefore, the overall renaturation rate associated with the repetitive c-ssDNAs will be the sum of rates corresponding to the incorrect contact and trap correct contact routes as described in **Eq. 13**. When the sequence complexity is close to the length of the c-ssDNAs, then the number of tcc and the associated slithering lengths will be much limited. This follows from the fact that the stability of incorrect contacts will be much lower than the trap correct contacts. As a result, the pathway via incorrect contacts will be the dominating one when *c* is close to *L* that is characterized by the scaling 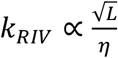. The number of tcc and the extent of slithering lengths will be much high when the sequence complexity is much lesser than the length of c-ssDNAs. Under such conditions, the pathway via trap correct contacts will be the dominating one which is characterized by the scaling 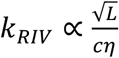 in line with the experimental observations [8]. Clear differences between one-step, two-step and three-step hybridization models are summarized in **Table 2**.

**TABLE 2.** Comparison of various hybridization models.

| Model | Summary | Predictions and limitations |
| --- | --- | --- |
| One-step model (Scheme I) | Hybridization occurs via one-step 3D diffusion controlled collision. | (1) This model predicts an inverse scaling of the hybridization rate with the length of DNA which contradicts the experimental findings that the hybridization rate scales with the length of DNA in a square root manner. (2) zipping time is ignored. Therefore, this model will work only for short DNAs. (3) Correctly predicts the viscosity dependence of the hybridization rate. |
| Two-step model (Scheme II) | Hybridization occurs via nucleation in the first step that is followed by zipping in the second step. | (1) This model predicts correct scaling of the hybridization rate with the length and complexity of DNA as long as the complexity is much lower than the length. However, when the complexity approaches the length, this model predicts inverse square root scaling of the hybridization rate with the length of DNA which is against the experimental observations. (2) Translational diffusion of DNA is ignored and the microscopic diffusion dynamics at the time of nucleation is used to explain the viscosity dependence of the hybridization rate. |
| Three-step model presented (Schemes III and IV) | Hybridization occurs via 3D diffusion mediated incorrect as well as trap-correct contacts in the first step, followed by nucleation via 1D slithering in the second step, followed by zipping in the third-step. In Scheme IV, slithering pathways are different for incorrect and trap-correct contacts mediated nucleation. | (1) Detailed contributions of trap-correct contacts arising from the repetitive DNA over the hybridization rate were not considered in Scheme III.<br>(2) Scheme IV, correctly predicts (a) the length and complexity dependence of the hybridization rate irrespective of the relative levels of complexity and length of the interacting DNA sequences. (b) Correctly predicts the viscosity dependence of the hybridization rate. |

### 4.1. Computational evidences for the three-step model

The single molecule coarse grained molecular dynamic trajectory of the hybridization of oxDNA showed a four-state mechanism which comprised [29] of random collisions between the complementary ssDNAs with oscillations in the systematic energy that is followed by nucleation, zippering and formation of duplex DNA. Here the oscillations in the systematic energy upon random collisions between c-ssDNAs represents the existence of several rounds of incorrect-contact-formations and dissociations of our model. The step corresponding to the initialization of hybridization represents the nucleation. Remarkably, the average zipping time seems to be almost independent on the electrostatic interactions arising at the DNA-DNA interface of c-ssDNAs which implies that the zippering is not a rate determining step especially for short c-ssDNA segments [29]. However, our theory predicts a linear scaling of the zipping time with the length of c-ssDNAs which means that the zipping time cannot be ignored when the length of c-ssDNAs is very large.

Coarse grained molecular dynamics simulation [33] of the hybridization of repetitive ATATATATAT oligomer revealed the existence of out-of-register shifted metastable states (partially hybridized DNA molecules at tcc with single strand overhangs of our model). The rate of formation of dsDNA from these metastable states seems to be twice faster than the formation rate from the corresponding fully dissociated c-ssDNAs [33]. This observation is in line with our theoretical prediction that the partially hybridized metastable states via tcc retain the c-ssDNAs in close vicinity for prolonged timescales that enhances the 1D slithering type search for the correct contacts to initiate the nucleation. This enhancement effect can be represented by the inequality condition *q* < *q_t_* in **Eq. 13**. However, our theory also predicted the retardation effects of such metastable trap configurations on the overall hybridization rate which can be represented by the inequality *k_r_* > *k_rt_* in **Eq. 13**. Insertion of a GC pair at the terminal or middle position of this poly AT oligomer, drastically reduced the number of out-of-register shifted metastable states leading to a two-state hybridization scheme [33]. This observation suggested that the ruggedness of the interaction energy landscape of the single stranded poly AT oligomers gets smoothened and funnelled upon insertion of a GC pair. That is to say, those out-of-state shifted metastable states are funnelled towards the stable GC contact-nucleus which indirectly reveals the presence of 1D slithering dynamics.

The possibility of bipedal directed walks of short fragments of DNA over a template DNA track have been demonstrated by several groups [43–47]. The directional dependent movement of these DNA walkers requires the input of an external energy. In this context, according to our three-step model, the correct contact formation across the c-ssDNAs is via a combination of 1D and 3D diffusion routes which is an unbiased random walk that is mainly driven by the background thermal energy.

Although the computational studies could reveal the finer details about the mechanism of DNA hybridization at the molecular level, our simplified lattice model is able to capture the overall dynamical aspects of hybridization phenomenon. Particularly, the three-step **Scheme IV** corresponding to the repetitive c-ssDNA along with **Eq. 13** can successfully explain how the overall hybridization rate scales with the viscosity of the reaction medium, and relative levels of lengths and sequence complexities of c-ssDNAs (particularly in the limit as the level of sequence complexity approaches the length) where most of the earlier theoretical models and computational studies failed.

### 4.2. Assumptions and limitations

Although the presented lattice model could recover several kinetic scaling laws associated with the DNA renaturation dynamics, there are several assumptions and limitations viz. 1) the rigidities of both c-ssDNAs and dsDNA are ignored. 2) same intermonomer distance for both c-ssDNA and dsDNA strands were assumed. 3) detailed base-paring and base stacking interactions were ignored. 4) formation of intra strand loops were ignored and (5) GC base compositions of the c-ssDNA were not considered.

Translation diffusion of c-ssDNAs and their interpenetration upon collision mainly decide the kinetic scaling laws rather than the fine details of base-pairing and base-stacking interactions. Therefore, these factors will not affect the main scaling results much. Variation in the GC composition of c-ssDNAs will influence the microscopic zipping rates k_+_ and k_−_ and subsequently the overall zipping rate *k_z_*. However, this will not affect the scaling of the zipping rate with the length of c-ssDNAs. Variation of the intermonomer distances of c-ssDNAs will not affect the obtained scaling laws since we measure the lengths in terms of number of monomer units (nt). For example, we consider only the number of nucleotides scanned by the c-ssDNAs over slithering events rather than the number of intermolecular distances. The rigidity of both c-ssDNA and dsDNA will be mainly compensated by the conformational fluctuations driven by the chain entropy [48]. Formation of intra strand loop structures will influence our results in two different ways viz. 1) significant fraction of c-ssDNA segments will not be exposed for the exploration of correct contacts which in turn increases the time required for nucleation. 2) upon forming correct contacts at the non-loop regions, zipping will be hindered by the presence of intra strand loops which in turn increases the overall zipping time. This means that additional time components will be added up to the overall nucleation an zipping times that will not change the overall kinetic scaling laws much.

## 5. Conclusion

We have developed a lattice model on the rate of hybridization of the complementary ssDNA (c-ssDNA) strands. These c-ssDNAs can be thought as loosely packed and spherical shaped nucleotide clusters and the collisions between these clusters are catered via the three-dimensional translational diffusion. Upon each collision, the base clusters of c-ssDNAs interpenetrate each other to form three different types of contacts among them viz. correct, incorrect and trap-correct contacts. Correct contacts are those with exact registry matches which can lead to nucleation and zipping of c-ssDNAs. Incorrect contacts are the mismatch contacts which are less stable compared to the trap correct contacts which can occur in the repetitive c-ssDNAs. Trap correct contacts possess exact registry match within the repeats. However, they are incorrect contacts in the view of the whole c-ssDNAs.

Trap correct contacts can form partial duplexes with single stranded overhangs. The nucleation rate (*k_N_*) is directly proportional to the average number of correct-contacts (<*n_cc_*>) formed when c-ssDNAs interpenetrate each other inside the reaction volume. Here the c-ssDNAs reach the reaction volume via translational diffusion with rate *k_t_*. This rate will be independent on the length of c-ssDNAs when both the c-ssDNAs are equal in size. As a result, the nucleation rate will be directly proportional to *k_t_* times the number of correct contacts. For short c-ssDNAs, the zipping times will much lower than the nucleation times. In such conditions, the overall renaturation rate (*k_R_*) will be directly proportional to *k_N_* and *L* as *k_R_* = *k_N_ L* ∝ *k_t_* <*n_cc_*> *L* which is the 3D diffusion model.

To understand the scaling properties of the average number of various types of contacts with respect to the length (*L*) of c-ssDNAs, sequence complexity (*c*), and reaction volume (*V*), we modelled the c-ssDNAs as a pair of self-avoiding random walks confined in a cubic lattice box which resembles the reaction volume (*V*). Our lattice model simulations suggested the scaling 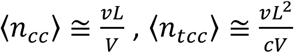 and 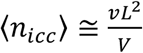 where the *v* = 1 is the monomer volume, 〈*n_Q_*〉 are the average number of correct (Q = cc), incorrect (Q = icc) and trap correct contacts (Q = tcc). Further numerical analysis and nonlinear least square fitting results suggested the scaling for the average radius of gyration of c-ssDNAs with their length as 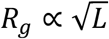. Since the reaction space will be approximately a sphere with radius equals to 2*R_g_*, one obtains the scaling for the nucleation rate with length of c-ssDNA as 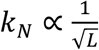 and one finally obtains 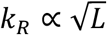 in line with the experimental observations. However, this expression works only for a nonrepetitive and short c-ssDNAs. When c-ssDNAs are repetitive with a complexity of *c* < *L*, then earlier models suggested the scaling 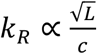. This scaling will break down when *c* = *L*. These observations clearly suggested the existence of at least two different pathways of renaturation viz. through incorrect contact and trap correct contact.

The trap correct contacts occurring between repetitive c-ssDNAs can lead to the formation of partial duplexes with single strand overhangs. These partial duplexes keep the c-ssDNAs in close vicinity for prolonged timescales which is essential for the extended 1D slithering that can speed up the searching for the correct contacts. Clearly, the extent of slithering dynamics will be inversely proportional to the sequence complexity *c*. When the complexity is close to the length of c-ssDNAs, then the pathway through incorrect contact will be the dominating one with minimal level of slithering. When the complexity is much lesser than the length of c-ssDNAs, then the pathway via the trap correct contact will be the dominating one.

